# Glycoprotein 3 of porcine reproductive and respiratory syndrome virus exhibits an unusual hairpin-like membrane topology

**DOI:** 10.1101/304337

**Authors:** Minze Zhang, Ludwig Krabben, Fangkun Wang, Michael Veit

## Abstract

The glycoprotein GP3 of the Arterivirus porcine reproductive and respiratory syndrome virus (PRRSV) consists of a cleaved signal peptide, a highly glycosylated domain, a short hydrophobic region and an unglycosylated C-terminal domain. GP3 is supposed to form a complex with GP2 and GP4 in virus particles, but secretion of GP3 from cells has also been reported.

We analyzed the membrane topology of GP3 from various PRRSV strains. A fraction of the protein is secreted from transfected cells; GP3 from PRRSV-1 strains to a greater extent than GP3 from PRRSV-2 strains. This secretion behavior is reversed after exchange of the variable C-terminal domain. A fluorescence protease protection assay shows that the C-terminus of GP3, fused to GFP, is resistant against proteolytic digestion in permeabilized cells. Furthermore, glycosylation sites inserted into the C-terminal part of GP3 are used. Both experiments indicate that the C-terminus of GP3 is translocated into the lumen of the endoplasmic reticulum. Deletion of the conserved hydrophobic region greatly enhances secretion of GP3 and fusion of this domain to GFP promotes membrane anchorage. Bioinformatics suggests that the hydrophobic region might form an amphipathic helix. Accordingly, exchanging only a few amino acids in its hydrophilic face prevents and in its hydrophobic face enhances secretion of GP3. Exchanging the latter amino acids in the context of the viral genome did not affect release of virions, but released particles were not infectious. In sum, GP3 exhibits an unusual hairpin-like membrane topology that might explain why a fraction of the protein is secreted.

**IMPORTANCE:** The porcine reproductive and respiratory syndrome virus (PRRSV) is the most important pathogen in the pork industry. It causes persistent infections that lead to reduced weight gain of piglets; highly pathogenic strains even kill 90% of an infected pig population. PRRSV cannot be eliminated from pig farms by vaccination due to the large amino acid variability between the existing strains, especially in the glycoproteins. Here we analyzed basic structural features of glycoprotein 3 (GP3) from various PRRSV strains. We show that the protein exhibits an unusual hairpin-like membrane topology; membrane anchoring might occur via an amphipathic helix. This rather weak membrane anchor explains why a fraction of the protein is secreted from cells. Interestingly, PRRSV-1 strains secrete more GP3 than PRRSV-2. We speculate that secreted GP3 might play a role during PRRSV infection of pigs; it might serve as a decoy to distract antibodies away from virus particles.

## INTRODUCTION

Arteriviruses are a family of enveloped positive stranded RNA viruses comprising the prototype member equine arteritis virus (EAV), lactate dehydrogenase-elevating virus (LDV) of mice and the porcine reproductive and respiratory syndrome virus (PRRSV), currently a major pathogen in the pork industry (1, 2). PRRSV was previously divided into two distinct genotypes termed “European” and “North American”, but because of the low nucleotide identity they are now classified as two genera, PRRSV-1 and PRRSV-2, respectively (3). The early isolates Lelystad virus (LV, PRRSV-1, (4, 5)) and VR-2332 (PRRSV-2, (4)) serve as respective prototype strains. Since their discovery, both genotypes have spread worldwide and diversified rapidly by mutation and recombination, including the occurrence of highly pathogenic variants in China ((6), related to PRRSV-2) and Eastern Europe ((7)), strain Lena, related to PRRSV-1). PRRSV cannot be eliminated from pig farms by vaccination due to the large variability between the existing strains. Especially the glycoproteins show strong antigenic drift and exhibit large variation (up to 50%) in their amino acid sequence (8, 9).

Arterivirus particles contain a multitude of membrane proteins, the disulphide-linked GP5/M dimer and the GP2/3/4 complex, the small and hydrophobic E-protein and the ORF5a protein (10). The glycoproteins GP2, GP3 and GP4 as well as the E-protein are essential for replication of both EAV and PRRSV. Deleting their genes individually from the viral genome does not prevent budding of virus-like particles from transfected cells, but the particles are not infectious (11, 12). More detailed experiments with EAV showed that if just one of the genes encoding GP2, GP3 or Gp4 were deleted the resulting particles do not contain the GP2/3/4 complex and reduced amounts of the E protein (11). It was therefore concluded that the GP2/3/4 complex is essential for virus entry, in the case of PRRSV probably due to its interaction with the key cellular receptor CD 163 (13–16).

Despite the large amount of sequence information on PRRSV genomes only limited information is available on the structure of their membrane proteins. GP2 and GP4 are typical type I transmembrane proteins that form an intermolecular disulfide linkage in the ER. GP3, the focus of this study, consists of an N-terminal signal peptide, a 180 amino acids long and highly glycosylated domain, a hydrophobic region (20 aa) and a variable unglycosylated C-terminal domain (50-60 aa). Some, but not all glycosylation sites are required for virus replication and loss of certain sites affect the antigenicity of GP3 (13, 17, 18). A multitude of linear antigenic regions were identified in GP3 proteins from both PRRSV-1 and PRSSV-2 strains, but antibodies directed against most epitopes do not neutralize viral infectivity (19–25). However, it was recently reported that PRRSV viruses which escaped from the antibody response in experimentally infected pigs contain mutations not only in its main glycoprotein GP5, but also in GP3 (26, 27). In transfected and infected cells GP3 is retained in the endoplasmic reticulum (ER), the viral budding site.

Inconsistent results have been published about whether GP3s are structural proteins of Arterivirus particles. Only for EAV it has been clearly shown, that GP3 is present in a disulfide-linked trimer with GP2/4 in virus particles. Uniquely, the disulfide linkages between GP3 and GP2/4 are not formed in the ER, but only after release of virus particles from infected cells (11, 28). In contrast, GP3 from LDV is only weakly associated with membranes and at least a fraction of the protein is secreted (29). Initial studies with GP3 from the PRRSV-2 strain IAF-Klop showed that it is secreted both from transfected and infected cells in a membrane-free form as disulphide-linked dimer carrying complex type carbohydrates (30). In contrast, a more recent study convincingly demonstrated that GP3 from the PRRSV-II FL-12 strain is incorporated into virus particles (31) as is GP3 from PRRSV-1 strain Lelystad (32). Since in the latter two studies it was not investigated whether a fraction of GP3 is secreted from transfected cells, we systematically examined whether this is the case for GP3 proteins from various PRRSV-1 and PRRSV-2 strains.

The GP3 protein seems to take a special position among the arterivirus glycoproteins; especially its membrane topology has not been resolved and remains speculative. For GP3 from EAV we showed that carbohydrates attached to overlapping glycosylation sites (NNTT) adjacent to the signal peptide prevents signal peptide cleavage. However, the uncleaved signal peptide does not act as a membrane anchor. Membrane attachment is caused by the hydrophobic C-terminus of GP3, which does not span the membrane, but rather attaches the protein peripherally to ER membranes (33, 34). In contrast, the signal peptide of GP3 from PRRSV (and of LDV) is cleaved despite the presence of a carbohydrate in its vicinity (35) indicating that there are differences in protein processing between the Arterivirus genera Rodartevirus (LDV and PRRSV) and Equartevirus (EAV).

Here we systematically investigated whether GP3 proteins from PRRSV-1 and PRRSV-2 strains are secreted from transfected cells. We also show that the C-terminal part of GP3 is translocated into the lumen of the ER. Membrane anchoring is achieved by a short hydrophobic region that might form an amphipathic helix.

## MATERIAL AND METHODS

### Cells

Cell lines CHO-K1 (Chinese hamster ovary cells, ATCC CCL-61), BHK-21 (baby hamster kidney cells, ATCC C13), 293T (Human embryonic kidney cells, ATCC CRL-3216), and MARC-145 (simian kidney epithelial cells derived from MA-104, ATCC CRL-6489) were maintained as adherent culture in Dulbecco’s Modified Eagle’s Medium (DMEM, PAN, Aidenbach, Germany) supplemented with 10% fetal calf serum (FCS) (Perbio, Bonn, Germany), 100U of penicillin per ml, and 100 mg of streptomycin per ml at 37°C in an atmosphere with 5% CO_2_ and 95% humidity.

### Plasmids and mutagenesis

The nucleotide sequences of open reading frame 3 of the following five strains of PRRSV was synthesized by Bio Basic Inc. (Markham Ontario, Canada): Lelystad virus (LV): low pathogenic PRRSV-1 prototype strain (36), GenBank accession number: M96262.2; VR-2332: low pathogenic PRRSV-2 prototype strain (37), accession number: U87392.3; IAF-Klop: PRRSV-2 Québec reference strain (38), accession number: AF003344; XH-GD: Chinese highly pathogenic PRRSV-2 strain, accession number EU624117.1 and Lena: highly pathogenic PRRSV-1 strain (7), accession number: JF802085.1. All GP3 genes were equipped during synthesis at the 3’end with a sequence encoding the HA-tag (amino acids YPYDVPDYA) plus a small linker (PV). The GP3 genes with HA-tag were subcloned into the plasmid pCMV-TNT (Promega, (Mannheim, Germany), containing T7 and CMV promotors) using KpnI and NotI restriction sites.

Using these plasmids as templates, site-directed mutagenesis was performed by overlap extension polymerase chain reaction (PCR) using standard molecular biology techniques to generate the following GP3-HA mutants: VR-2332 GP3 with point mutations L227N, A246N and Lelystad (LV) GP3 with point mutations F228N and V255S to introduce N-glycosylation sites, VR-2332 GP3 with point mutations N195Q and Lelystad virus GP3 with point mutations N194Q to delete a N-glycosylation site, VR-2332 GP3 with mutations in the predicted amphipathic helix: GP3-2A (L194A W198A), GP3-3A (L184A F187A W191A) and GP3-5A (L184A F187A W191A L194A W198A),. XH-GD GP3 with mutations in the predicted amphipathic helix: GP3-3H (N195S S197L W198L), GP3-4H (R185L P186L S189F S190F) and GP3-7H (R185L P186L S189F S190F N195S S197L W198L). These mutations in the predicted amphipathic helix were also performed in the context of mutant GP3A246N of VR-2332: GP3-3H (N195S S197L W198L), GP3-4H (R185L P186L S189F S190F) and GP3-7H (R185L P186L S189F S190F N195S S197L W198L).

To delete parts from the C-terminus of GP3 of VR-2332, nucleotides encoding the amino acids sequence 1-229 (GP3-△C2) or 1-177 (GP3-△C1) of GP3 were amplified by PCR using forward primers equipped with KpnI site or reversed primers encoding the HA-tag (amino acids YPYDVPDYA) including the small linker (PV) and equipped with a NotI site. Mutant GP3-△C2 lacks the between strains variable region of GP3 and GP3-△C1 also the hydrophobic region. The resulting sequences were subcloned into the plasmid pCMV-TNT using the same enzymes. This strategy was also used for construction of mutant GP3-△C1 of

Lelystad virus (LV) that contains the amino acids sequence 1-171 and mutants GP3-△C that contains the amino acids residues 1-203 of GP3 of Lelystad virus (LV) or 1-204 of GP3 of XH-GD. GP5-HA has been described (39).

The two chimeras LV-XH-GD and XH-GD-LV were constructed using the GP3-wt plasmids of the PRRSV strains Lelystad virus (LV) and XH-GD as a template for overlap extension polymerase chain reaction (PCR) using standard molecular biology techniques. The chimera LV-XH-GD contains the amino acids 1-203 of GP3 of LV (including hydrophobic region) plus the amino acids 205-254 of GP3 of XH-GD (complete C-terminus excluding hydrophobic region). The chimera XH-GD-LV contains the amino acids 1-204 of GP3 of XH-GD (including hydrophobic region) plus the amino acids 204-265 of GP3 of LV (complete C-terminus excluding hydrophobic region). Both chimeras contain the HA-tag (amino acids YPYDVPDYA) and the small linker (PV) as described above.

GP3-YFPs were constructed using plasmids GP3-HA (wild type) of the five PRRSV strains described above as a template. The full length GP3 gene was amplified by PCR using primers equipped with BamHI or KpnI sites and subcloned into pEYFP-N1 (Clontech, Saint-Germain-en-Laye, France) using the same enzymes. This procedure inserts the linker sequence RPVATM between GP3 and YFP, which is derived from the cloning site of the vector. GP4-YFP was constructed as described (33) using the same strategy.

To fuse different regions from the C-terminus of GP3 of VR-2332 to the C-terminus of GFP, nucleotides encoding the amino acids sequences 178-254 (encoding the complete C-terminus including the hydrophobic region), 178-229 (the between strains variable region is not present) or 181-200 (hydrophobic region only) were amplified by PCR (primers equipped with BglII / Hind III or BamHI sites) and cloned to the corresponding sites in in pEGFP-C1 (Clontech, Saint-Germain-en-Laye, France). The resulting constructs are called GFP-HR1 (containing amino acids 181-200), GFP-HR2 (178-229) and GFP-HR3 (178-254), respectively. The cloning procedure adds the following amino acids between GFP and the GP3 sequence: GFP-HR1: RPDSDLEL; GFP-HR2 and GFP-HR3: SGL. The GP3 gene of all plasmids was sequenced (LGC Genomics, Berlin, Germany) before use in experiments.

### Transfection and analyzing intracellular and secreted GP3

Cells in 6-well plates were transfected with 2.5 μg plasmid DNA using Lipofectamine^®^ 3000 (Thermo Fisher Scientific, Carlsbad, United States) as described by the manufacturer. Twenty-four hours after transfection of cells, cellular supernatants were removed and cleared from cell debris by low speed centrifugation (3000xg, 10 min, 4°C). Proteins in the supernatant were precipitated with trichloroacetic acid (TCA, 10% final concentration) and pelleted (15000g, 30min, 4°C). Pellets were washed three times with 100% ethanol, dried and resuspended in reducing SDS-PAGE buffer.

To prepare intracellular GP3, cells were washed with PBS, detached from the dish with trypsin-EDTA (PAN Biotech, Aidenbach, Germany), pelleted, washed twice with phosphate-buffered saline (PBS, 137mM NaCl, 2.7mM KCl, 10mM Na2HPO4, 1.8mM KH2PO4, pH 7.4), resuspended in glycoprotein denaturing buffer (0.5% SDS, 40 mM DTT) and boiled for 10 min at 100°C.

To analyze secretion of GP3 in virus-infected cells, MARC-145 cells were infected with XH-GD or Lelystad virus (LV) at a moi of ~ 1 four hours after transfection and samples (cellular lysates and supernatants) were processed 24 hours after transfection as described above.

To analyze glycosylation samples were digested with Peptide-N-Glycosidase (PNGase F, 2.5-5units/μL, 4 h at 37°C) or endo-beta-N-acetyl-glucosaminidase (Endo H, 2.5-5units/μL, 1 h at 37°C) according to the manufacturer’s instruction (New England Biolabs, Frankfurt am Main, Germany). For limited PNGase F digestion, a serial 10-fold dilution of PNGase F (starting with 1 unit/μL) was prepared for incubation with the substrate. Deglycosylated or untreated samples were supplemented with reducing (in one experiment non-reducing) SDS-PAGE loading buffer and subjected to SDS-PAGE and Western blot.

### Membrane fractionation of transfected cells

Two different membrane fractionation assays were performed. The first method separates microsomes and cytosol (Fig. 5), the second firstly prepares microsomes and then separates their luminal (soluble) and membranous content (Fig. 4). Twenty-four hours after transfection of 293T cells cellular supernatants were discarded. Cells were washed twice with ice-cold PBS and treated with digitonin (30μM) for permeabilization for 25 minutes on ice. Nuclei and cell debris were removed by low speed centrifugation (700xg, 3min, Eppendorf centrifuge). Microsomes (ER and Golgi) were recovered from the supernatant by high speed centrifugation (100000g, 1h, 4°C, 60 min., Beckman TL100 centrifuge, rotor TLA100.3). The supernatant of this centrifugation contains the cytosolic proteins, which were precipitated with TCA as described above.

Pellets of microsomes were either resuspended in reducing SDS-PAGE buffer (method 1) or were opened (method 2) using ultrasonication (Misonix XL-2000). Membranous (membrane-associated protein) and soluble fractions (soluble protein in the lumen of ER) were separated by centrifugation (100000g, 1h, 4°C). The membranous pellet was washed once in PBS and resuspended in reducing SDS-PAGE buffer. Soluble proteins were precipitated with TCA as described above.

### SDS-PAGE and Western Blot

After sodium dodecyl sulfate-polyacrylamide gel electrophoresis (SDS-PAGE) using 15% or 12% polyacrylamide gels were blotted onto polyvinylidene difluoride (PVDF) membrane (GE Healthcare, Freiburg im Breisgau, Germany). After blocking of membranes (blocking solution: 5% skim milk powder in PBS with 0.1% Tween-20 (PBST)) for 1 h at room temperature, antibodies in blocking solution were applied for overnight at 4°C: rabbit-anti-HA tag antibody (ab9110, Abcam, Cambridge, UK, 1:10.000) was used to detect GP3 with HA tag and GP5 with HA tag, Mouse-anti-N antibody (13E2) (reactive against PRRSV-1 strains, 1:1000), Mouse monoclonal anti-GP5 antibody (reactive against PRRSV-2 strains, 1: 1000). After washing (3×10 min with PBST), suitable horseradish peroxidase-coupled secondary antibody (anti-rabbit or anti-mouse, Sigma-Aldrich, Taufkirchen, Germany, 1:5.000) was applied for 1 hour at room temperature. After washing, signals were detected by chemiluminescence using the ECLplus reagent (Pierce/Thermo, Bonn, Germany) and a Fusion SL camera system (Peqlab, Erlangen, Germany).

### Immunofluorescence Assay

Transfected CHO-K1 or infected MARC-145 cells grown in 6-well plates (Sarstedt, Nümbrecht, Germany) were fixed with paraformaldehyde (4% in phosphate-buffered saline (PBS)) for 20 min at 4°C, washed twice with PBS, permeabilized with 0.5% Triton in PBS for 10 min at room temperature, and washed again twice with PBS. After blocking (blocking solution: 3% bovine serum albumin (BSA) in PBS with 0.1% Tween-20 (PBST)) for 30 min at room temperature the cells were incubated with a mouse monoclonal anti-GP5 antibody diluted in blocking solution (reactive against PRRSV-2 strains, 1: 100) at room temperature for 1 hour, then washed three times with PBS, incubated with secondary antibody (Alex Fluor 568 goat anti-mouse IgG(H+L), Invitrogen, Darmstadt, Germany, 1:1000). Pictures were recorded using a ZEISS Axio Vert. A1 inverse epifluorescence microscope.

### Fluorescence protease protection assay

The fluorescence protease protection assay was performed as described (33). Briefly, CHO-K1 cells seeded on 6-well plates were transfected with plasmids GP3-YFP of all PRRSV strains or GP4-YFP of EAV as described above. Twenty-four hours after transfection cells were treated with digitonin (30μM) for 1min, and then proteinase K (50ng/ml, Sigma-Aldrich) was added. Pictures were recorded using a ZEISS Axio Vert. A1 inverse epifluorescence microscope.

### Mutagenesis and reverse genetics with full-length PRRSV cDNA clone

The full-length infectious PRRSV cDNA clone pOKXH-GD (a kind gift of Prof. Guihong Zhang, South China Agricultural University) was constructed from the XH-GD PRRSV strain (GenBank accession no. EU624117) and vector pOKq (40). The entire genome of PRRSV is present in four plasmids: the plasmid pOK-A contains a CMV promotor plus the PRRSV nucleotide sequences 1 to 6465 (fragment A), pOK-B nucleotide sequences 6466 to 9301 (fragment B), pOK-C 9302 to 12586 (fragment C), and pOK-D nucleotide sequences 12587 to 15345 plus a terminal BGH RNA transcription terminator sequence (fragment D).

Site-directed mutagenesis using overlap extension polymerase chain reaction (PCR) was performed with the pOK-D plasmid to introduce the same mutations in the hydrophobic region as in GP3 of VR-2332. GP3-2A contains the mutations L194A plus W198A, pGP3-3A the mutations L184A + F187A + W191A and pGP3-5A all five mutations, L184A + F187A + W191A + L194A + W198A. It was not possible to create the mentioned mutations in GP3 without affecting the coding sequence of the overlapping GP4 gene: the mutation L184A in GP3 changed A2G in GP4, mutation F187A changed F5C; mutation W191A changed L9C; mutation L194A changed F12C and mutation W198A changed V16G. The corresponding amino acids in GP4 are all located in the middle of the signal peptide, which is cleaved in EAV after amino acid 21 (10) and predicted by SignalP (http://www.cbs.dtu.dk/services/SignalP/) to be cleaved in PRRSV XH-GD after amino acid 22. Since this part of the signal peptide tolerates amino acid substitutions (Fig. 3A) and since SignalP also predicts in all mutants a cleaved signal peptide, the mutations are unlikely to affect the functionality of GP4.

The GP3 genes of the pOK-D plasmids containing the desired mutations were completely sequenced to exclude unspecific second site mutations introduced by PCR. The C fragment was cut from pOK-C plasmid by digestion with BamHI and HindIII and ligated to the pOK-D plasmid digested with the same enzymes. The B fragment was cut from pOK-B plasmid by digestion with BamHI and AfIII and ligated into the resulting pOK-CD plasmid digested with the same enzymes. Finally, the A fragment was cut from the pOK-A plasmid by digestion with AfIII and NotI and ligated with the resulting pOK-BCD plasmid digested with the same enzymes. The complete plasmid was also constructed from wild type pOK-D in parallel. After each ligation the resulting plasmids were digested with four different restriction enzymes. The resulting band pattern was identical between the three mutants and wild-type indicating that the procedure did not introduce deletions or insertions into the PRRSV cDNA.

The plasmids (2.5 μg) containing the complete PRRSV genome were transfected into 80% confluent CHO-K1 or BHK-21 cells grown in 6-well plates using Lipofectamine^®^ 3000 (Thermo Fisher Scientific, Carlsbad, United States) as described by the manufacturer. After 48 hours the supernatant was removed, and cells were subjected to immunofluorescence assay using anti-GP5 specific antibody. The supernatant was cleared by low speed centrifugation and 500μl was used to infect (PBS washed) MARC-145 cells grown to 70-80% confluence on 6-well plates. After incubation for 1 h at 37 °C, the inoculums were removed, cells were washed twice with PBS, and cells were incubated in culture medium (DMEM with 2% FCS) for 48 h. Cells were then subjected to immunofluorescence assay using anti-GP5 specific monoclonal antibody.

### RNA isolation and RNA quantification by real time PCR

To calculate the amount of viral genomic RNA (as a measure of total viral particles) released into the supernatant of transfected or infected cells, quantitative real time PCR was performed. Supernatants (1-2ml) were cleared by low speed centrifugation (10min, 5000g) and 30μl RNA is extracted from 400μl supernatant from either transfected or infected cells using PureLink viral RNA/DNA Mini Kit (Invitrogen, Carlsbad, United States). 8μl RNA is then supplemented with 1μl 10xbuffer and 1μl RNase-free DNase and incubated at 37° for 45min to remove genomic DNA. The reaction is stopped by adding 1μl DNase stop solution. Samples are then reverse transcribed into cDNA using High Capacity cDNA reverse transcription kit (Applied Biosystems, Carlsbad, United States). The cDNA yield was 800-900ng/μl in a volume of 21μl. Samples are diluted to 300ng/μl for qRT-PCR.

A standard curve was generated by serial dilution (10^8^-10^1^ copies/μl) of the full-length cDNA clone pGP3-wt plasmid. The diluted plasmids and the cDNA from samples were used for qRT-PCR, which was performed using the StepOnePlus real-time PCR system (Applied Biosystems) with SYBR green as fluorophore. The primers amplified a 180 bp fragment (427bp -606bp) from the GP3 gene. Gene copy numbers were calculated with StepOne Software v2.3 (Applied Biosystems). Results from two independent transfections or infections are shown as number of viral RNA (mean including standard deviation) released into the supernatant of transfected or infected 6-well cell culture plates (~5×105 cells).

## RESULTS

### A fraction of GP3 is secreted from transfected cells, but the amount varies between GP3s from PRRSV-1 and PRRSV-2 strains

To systematically investigate secretion of GP3 we expressed the ORF3 gene from the following five PRRSV strains in BHK-21 cells: from a low-pathogenic (Lelystad) and a highly pathogenic (Lena) “European” PRRSV-1 strain and two low-pathogenic (VR-2332, IAF-Klop) and one highly pathogenic (XH-GD) “North American” PRRSV-2 strain. All constructs were fused at their C-termini to an HA-tag for subsequent antibody detection. Western-blotting of cellular lysates and supernatants showed that GP3 from all five PRRSV strains is secreted, but to a different extent (Fig. 1A). Whereas the band intensity of intracellular GP3 is similar for all five GP3 proteins, the extracellular GP3 bands from the PRRSV-1 strains Lelystad and Lena are much more prominent than the corresponding band from the three PRRSV-2 strains. Similar results were obtained using transfection of CHO-K1 cells (data not shown). Quantification of band intensities (lysate and supernatant) revealed that approximately 20-30% of total GP3 from PRRSV-1 strains were released, whereas ~5-10% of GP3 from PRRSV-2 strains was secreted.

**FIG.1. A.**
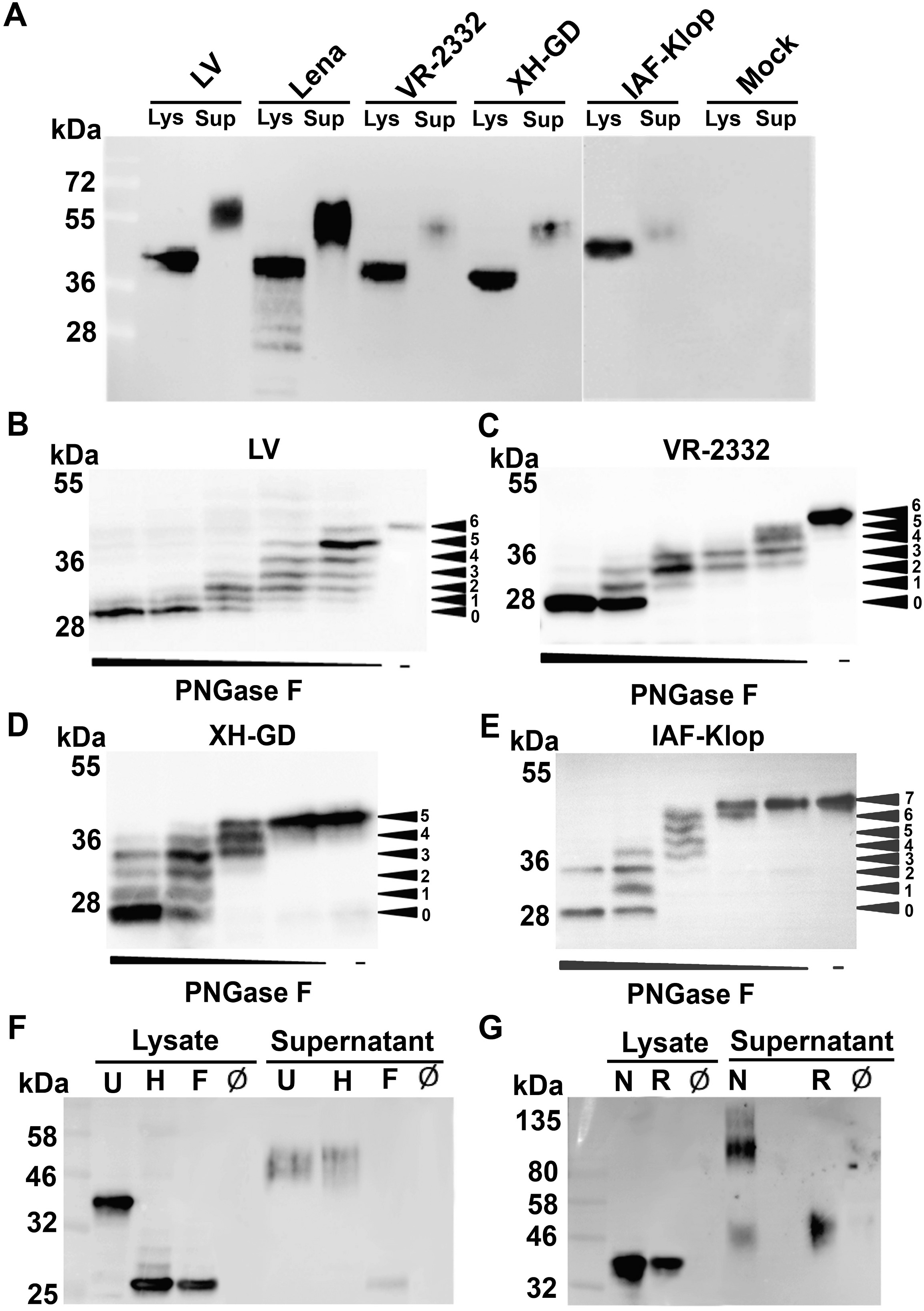
fraction of GP3 is secreted from transfected cells. (A) GP3 (equipped with a C-terminal HA-tag) from five PRRSV-1 and PRRSV-2 strains was expressed in BHK-21 cell. 24 hours after transfection the supernatant was removed, cleared and proteins were precipitated with TCA. Cells were washed, trypsinized, pelleted and lysed in denaturing buffer. Samples were subjected to reducing SDS-PAGE and Western blotting with anti-HA antibodies. To compare the amount of secreted GP3 between strains, different aliquots of the cell lysate were loaded onto the gel to achieve the same band intensity for intracellular GP3. The aliquot of the supernatant was adjusted accordingly for each GP3. Since ~ 5% (1-12.5%) of the total lysate, but much more (17.5-100%) of the supernatant was loaded onto the gel, the amount of secreted GP3 is overrepresented in this blot. Lelystad virus (LV): PRRSV-1 prototype strain, VR-2332: PRRSV-2 prototype strain; IAF-Klop: PRRSV-2 strain; XH-GD: highly pathogenic Chinese PRRSV-2 strain and Lena: highly pathogenic PRRSV-1 strain. Mock are untransfected cells. (B-E) GP3-wt from Lelystad (B), VR-2332 (C), XH-GD (D) and from IAF-Klop virus (E) were expressed in BHK-21 cells, cell lysates were not digested (-) or digested with serial 10-fold dilutions of PNGase F prior to Western blotting with anti-HA antibodies. This causes a ladder-like appearance of bands that allows counting of the total number of carbohydrates linked to GP3 (arrowheads). (F) GP3-wt from VR-2332 was expressed in BHK-21 cells, cell lysates and supernatants were not digested (U) or digested with PNGase F (F) or Endo H (H) prior to Western blotting with anti-HA antibodies. (G) GP3-wt from VR-2332 was expressed in BHK-21 cells, cell lysates and supernatants were subjected to reducing (R) or non-reducing (N) SDS-PAGE and Western blotting. The left lanes show molecular weight markers with the indicated sizes in kDa. Ø indicates untransfected cells.

Whereas the SDS-PAGE mobilities of the intracellular forms of GP3 from the strains Lelystad, Lena and VR-2332 are identical (~42 kDa), GP3 from XH-GD exhibits a slightly lower (~39 kDa) and GP3 from the IAF-Klop strain a clearly higher molecular weight (~45 kDa). Although the number of amino acids differ in GP3s of the strains investigated here (254 (including signal peptide) in PRRSV-2 strains, 249 in Lena and 265 in Lelystad), the calculated molecular weights are almost identical (30 – 32 kDa). However, GP3s vary in the number of predicted N-glycosylation sites; seven in Lelystad, Lena and VR-2332, six in XH-GD, but eight in IAF-Klop (see Fig. 3A for localization of glycosylation sites). To investigate how many glycosylation sites are used we carried out a limited digestion of cell lysates with PNGase F prior to western blotting. Counting of bands revealed that GP3 from Lelystad and VR-2332 contain six, GP3 from XH-GD five and GP3 from IAF-Klop seven oligosaccharides (Fig. 1B-E). Thus, in each of the GP3s all (except the one in the hydrophobic region, see Fig. 6) predicted glycosylation sites are used which explains the different SDS-PAGE mobilities of the glycosylated proteins.

GP3 present in the supernatant displayed a clearly higher molecular weight than intracellular GP3 and a smear band pattern often observed for glycoproteins subjected to terminal and heterogeneous glycosylation in the medial and trans-Golgi. To prove that GP3 was secreted via the exocytic pathway, intracellular and secreted GP3 from VR-2332 were digested prior to SDS-PAGE with PNGase F (that cleaves all types of N-linked carbohydrates) or Endo-H (which cleaves only high-mannose type carbohydrates typical for ER-localized proteins). The carbohydrates from intracellular GP3 are completely sensitive against both enzymes confirming that GP3 is retained in the ER or a cis-Golgi compartment (30, 41). In contrast, GP3 present in the supernatant is completely resistant against Endo-H digestion indicating that all its carbohydrates are terminally glycosylated (Fig. 1F). The much higher molecular weight of secreted versus intracellular GP3 is thus due to addition of several N-acetyl-glucosamine, galactose and sialic acid moieties to each of the glycosylation sites in the medial and trans Golgi. Note also that intracellular and secreted GP3 have an identical molecular weight after deglycoslyation with PNGase F indicating that the secreted form is not a result of proteolytic digestion, as has been reported for the secreted fraction of other viral spike proteins (42). Cleavage of GP3 just upstream of the hydrophobic region would remove 80 amino acids thereby yielding a product with a much higher SDS-PAGE mobility.

To assess the oligomeric nature of GP3 we subjected intracellular and secreted forms of GP3 from PRRSV-2 strain VR-2332 to non-reducing SDS-PAGE. Whereas intracellular GP3 revealed no size difference in the presence or absence of DTT, the majority of secreted GP3 showed a much higher molecular weight under non-reducing conditions, most likely representing a disulfide-linked dimer (Fig. 1G). Identical results were obtained if GP3 from the PRRSV-1 strain Lelystad was digested with glycosidases or subjected to non-reducing SDS-PAGE (data not shown). Likewise, GP3 of IAF-Klop was also reported to be secreted as an Endo-H resistant, disulfide-linked dimer (30).

### GP3 is also secreted in the context of a virus infection

One might argue that secretion of GP3 is an artifact of the expression system and the protein is not released from virus-infected cells, for example because it binds to the transmembrane proteins GP2 and/or GP4. To investigate this we transfected MARC-145 cells with GP3 from either XH-GD or from Lelystad virus, and infected four hours later with the corresponding virus. Cell lysates and supernatants (prepared 24 hours after transfection) were then tested with anti HA-antibodies to detect GP3 and with anti-GP5 and anti-N antibodies to monitor infection with XH-GD and Lelystad virus, respectively. The western blot shows that virus infection does neither change expression levels of GP3 inside cells nor its molecular weight (Fig. 2). In addition, GP3 was also detected in (almost) similar amounts in the supernatant of virus-infected cells. However, we cannot exclude that fusion of the HA-tag to GP3 prevents binding to GP2 and/or GP4, but for the IAF-Klop strain it was clearly shown that native GP3 is also secreted in a soluble, membrane-free form from virus-infected cells (30).

**FIG. 2.**
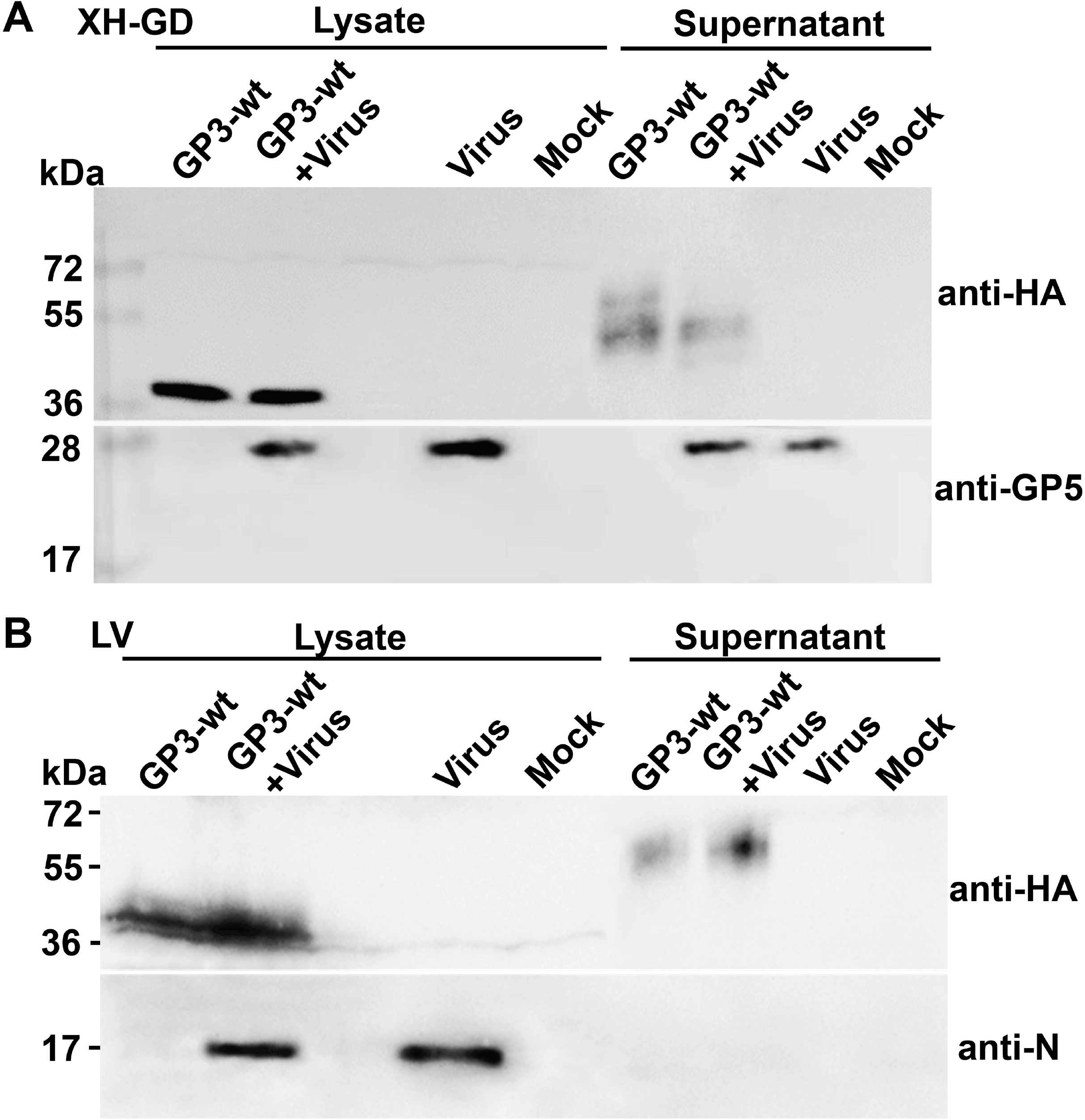
GP3 is also secreted from virus infected cells. (A) and (B) MARC-145 cells were transfected with GP3-wt of XH-GD (A) or Lelystad virus (LV) (B) and 4 hours later infected or not infected with the corresponding virus at an moi of ~1. 24 hours after transfection, when some cytopathic effects appeared, supernatants and cells were processed as described in figure legend 1. Samples were subjected to reducing SDS-PAGE and Western blotting with anti-HA (to detect GP3-HA), anti-GP5 (for XH-GD) and anti-N (for LV) antibodies. The left lanes show molecular weight markers with the indicated sizes in kDa. “Virus” indicates infected, but untransfected cells; “Mock” untransfected, non-infected cells. Note that the glycosylation pattern of GP3 slightly differs between transfected and infected cells.

### The C-terminus of GP3 is translocated into the lumen of the ER

A hydrophobic plot of GP3 revealed two hydrophobic regions; one is the N-terminal signal peptide, which we have shown to be cleaved in transfected cells (41), the second is an internal ~20 amino acid long hydrophobic region (HR). Since GP3 is a highly variable protein we also defined its conserved and variable parts. 619 ORF3 nucleotide sequences from both PRRSV-1 (162) and PRRSV-2 (457) strains present in the NCBI-database were translated into the corresponding amino acid sequences. From these data a consensus sequence for GP3 of both PRRSV-1 and PRRSV-2 containing the most abundant amino acid at each position was compiled and the percent conservation at each position was plotted against the amino acid number (Fig. 3A). The plot shows that the signal peptide (especially in its C-terminal part which contains the cleavage site) and the C-terminal part downstream of the hydrophobic region are the most variable regions of GP3. In contrast, the hydrophobic region is highly conserved. The domain between the signal peptide and the hydrophobic region is fairly conserved; completely non-variable parts are interrupted by one or a few variable positions. Features which mainly determine the structure of GP3, such as six (possibly disulfide-bond forming) cysteine residues and also seven sites for N-linked glycosylation (at position 42, 50, 131, 152, 160 and 195 in VR-2332) are completely conserved in this region. Another highly conserved glycosylation site is present adjacent to the signal peptide, but its position varies between PRRSV-1 (amino acid 27) and PRSSV-2 strains (amino acid 29). No glycosylation sites or cysteine residues are present in the variable C-terminal part of GP3, suggesting that this region might exhibit a flexible structure.

**FIG. 3.**
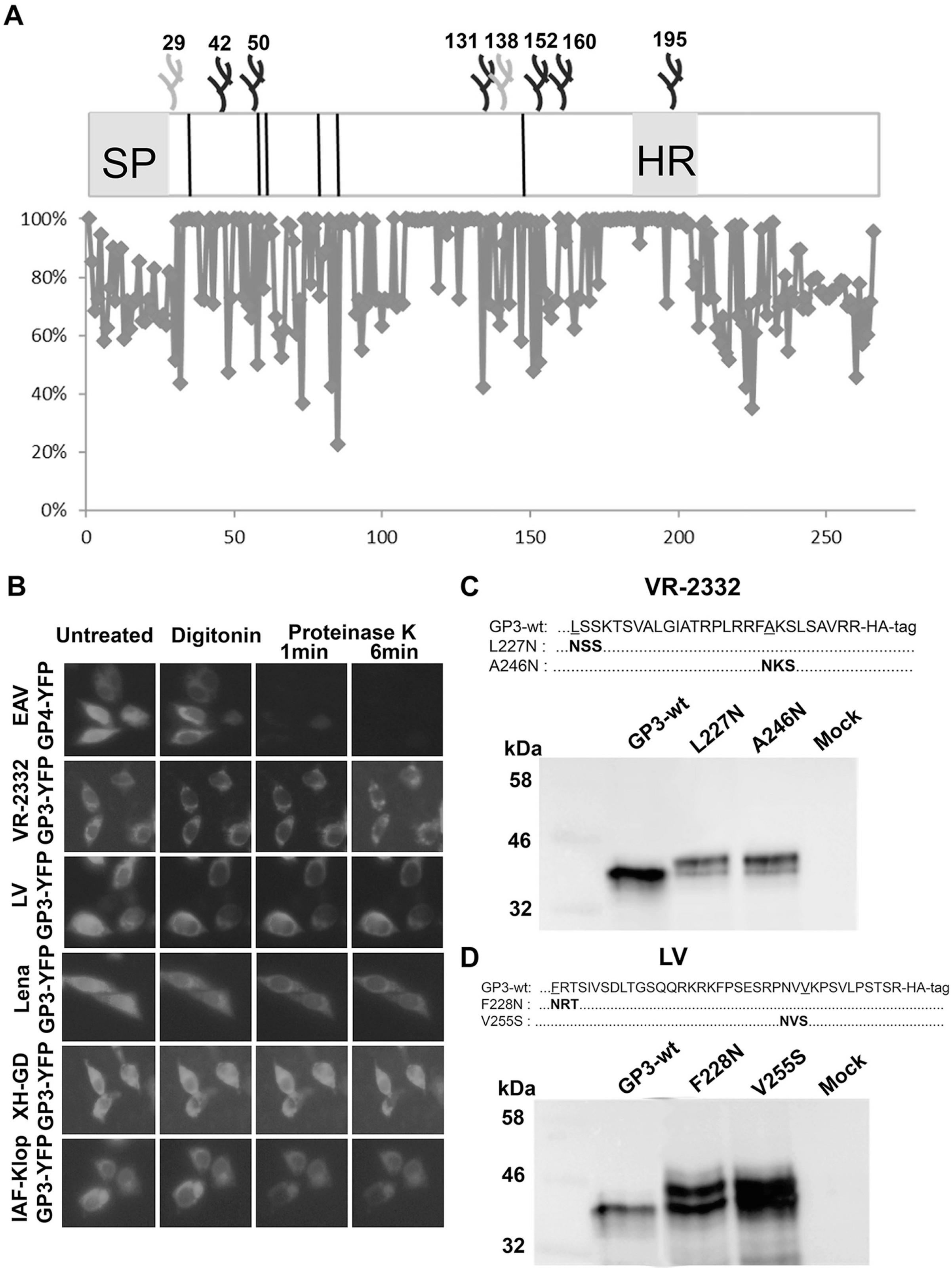
The C-terminus of GP3 is translocated into the lumen of the ER
(A) Primary structure of GP3 with signal peptide (SP), hydrophobic region (HR), six conserved cysteines (lines) and glycosylation sites (branches, numbering of sites is for PRRSV-2 strains), The position of the first site differs between PRRSV-1 and PRRSV-2 strains, glycosylation site 138 is an additional site present in GP3 from the IAF-Klop strain, GP3 from XH-GD lacks the site at position 152. The graph shows the percent conservation (Y-axis) of amino acids at each position (X-axis) of a consensus sequence compiled from all PRRSV-1 and PRRSV-2 GP3 sequences present in the database.
(B) CHO-K1 cells expressing GP3-YFP from 5 different PRRSV strains and (as a control) a type I transmembrane protein (GP4-YFP from equine arteritis virus (EAV)) were treated with digitonin for 1 min and with proteinase K for 1 and 6 min. After each time point, the same microscopic field was recorded with an epifluorescence microscope.
(C) and (D) Two additional glycosylation sites were inserted into the C-terminal part of GP3 from VR-2332 (C) or Lelystad virus (LV) (D) and constructs were expressed in BHK-cells. 24 hours after transfection cells were lysed and samples subjected to SDS-PAGE and Western blotting with anti-HA antibody. The C-terminal sequences of GP3 (amino acids 227-254 in VR-2332, 228-265 in LV) are shown above the blot. Bold letters indicate the N-glycosylation site inserted by exchange of one amino acid (underlined). The SDS-PAGE mobility of molecular weight makers are indicated at the left side of the blot. Mock: untransfected cells.

Considering the basic features of GP3 (Fig. 3A) two membrane topologies are possible: It might be a classical type I membrane protein with an N-terminal luminal glycosylated domain, the hydrophobic region is spanning the membrane and the C-terminus is a cytoplasmic tail. Alternatively, GP3 might form a loop-like structure; both the N-and C-terminus of the protein are present in the lumen of the ER and the hydrophobic region peripherally anchors GP3 to the membrane. We performed two different experiments to determine the membrane topology of GP3. First we used a fluorescence protease protection assay to investigate whether the C-terminus of GP3 is translocated into the ER lumen. The complete GP3 sequence of five PRRSV strains was fused at its C-terminus to YFP. The fluorophore will be protected from proteases added to perforated cells when it protrudes into the ER lumen, but not when exposed to the cytosol. As a control for a typical type I membrane protein with a C-terminus exposed to the cytoplasm, we used GP4 from equine arteritis virus (EAV) fused in a similar manner to YFP (33).

Fluorescence microscopy of transfected CHO-K1 cells showed comparable reticular ER staining for GP4-YFP and GP3-YFP of the five different PRRSV strains. Upon addition of the mild detergent digitonin, which solubilizes the plasma membrane, but not internal membranes, the fluorescence emitted from GP3-YFP and GP4-YFP was not diminished. However, only one minute after addition of Proteinase K, the same microscopic image section from cells expressing GP4-YFP did not show any fluorescence, indicating that YFP was digested. In contrast, even after six minutes of protease treatment, the fluorescence from GP3-YFP expressing cells was not even reduced (Fig. 3B). We conclude that the C-terminus of GP3 from PRRSV-1 and PRRSV-2 strains is exposed to the lumen of the ER.

To confirm this we inserted sites for N-linked glycosylation into the C-terminal part of GP3 from PRRSV-1 (Lelystad) and PRRSV-2 (VR-2332) strains. Since the active center of the oligosaccharyltransferase is exposed to the lumen of the ER, it can attach carbohydrates only to sites present in the same compartment (43). To minimize possible effects of the mutation on the structure of GP3 we exchange only one amino acid to create an N-glycosylation site at position 227 (mutation L to N) and 246 (A to N) of GP3 from VR-2332 and at position 228 (F to N) and 255 (V to S) of GP3 from Lelystad.

Western-blotting of cellular lysates showed one band for wildtype GP3, but two bands for all four mutants (Fig. 3C +D). The lower band ran at the same height as the corresponding wildtype GP3, whereas the other band showed a size increase consistent with the attachment of an additional glycan. In both mutants of GP3 from VR-2332 the upper band account for ~80% of total GP3, whereas in GP3 of Lelystad both fractions constitute ~ 50%. In principle, there are two possible explanations why the introduced sites are not stoichiometrically glycosylated: (i) The C-terminus might not be translocated into the ER lumen in every synthesized GP3 molecule or (ii) C-terminal glycosylation sites might be occasionally skipped because ongoing protein folding hides them from the oligosaccharyltransferase as a quantitative study of the mouse N-glycoproteome suggested (44). The latter possibility is more likely since YFP attached to the C-terminus of GP3 was completely resistant against proteolytic digestion in the fluorescence protection assay (Fig. 3B).

### A hydrophobic region in the C-terminal part of GP3 is the membrane anchor

Having established that the C-terminus of GP3 is translocated into the lumen of the ER, we next asked whether the hydrophobic region serves as a membrane anchor. We deleted two regions from the C-terminus of GP3 from VR-2332. The mutant GP3-∆C2 contains amino acids 1-229; the variable C-terminal region is thereby removed, but the hydrophobic region is still present. In GP3-∆C1 (containing amino acids 1-177) the hydrophobic region was also deleted (Fig. 4A).

**FIG. 4.**
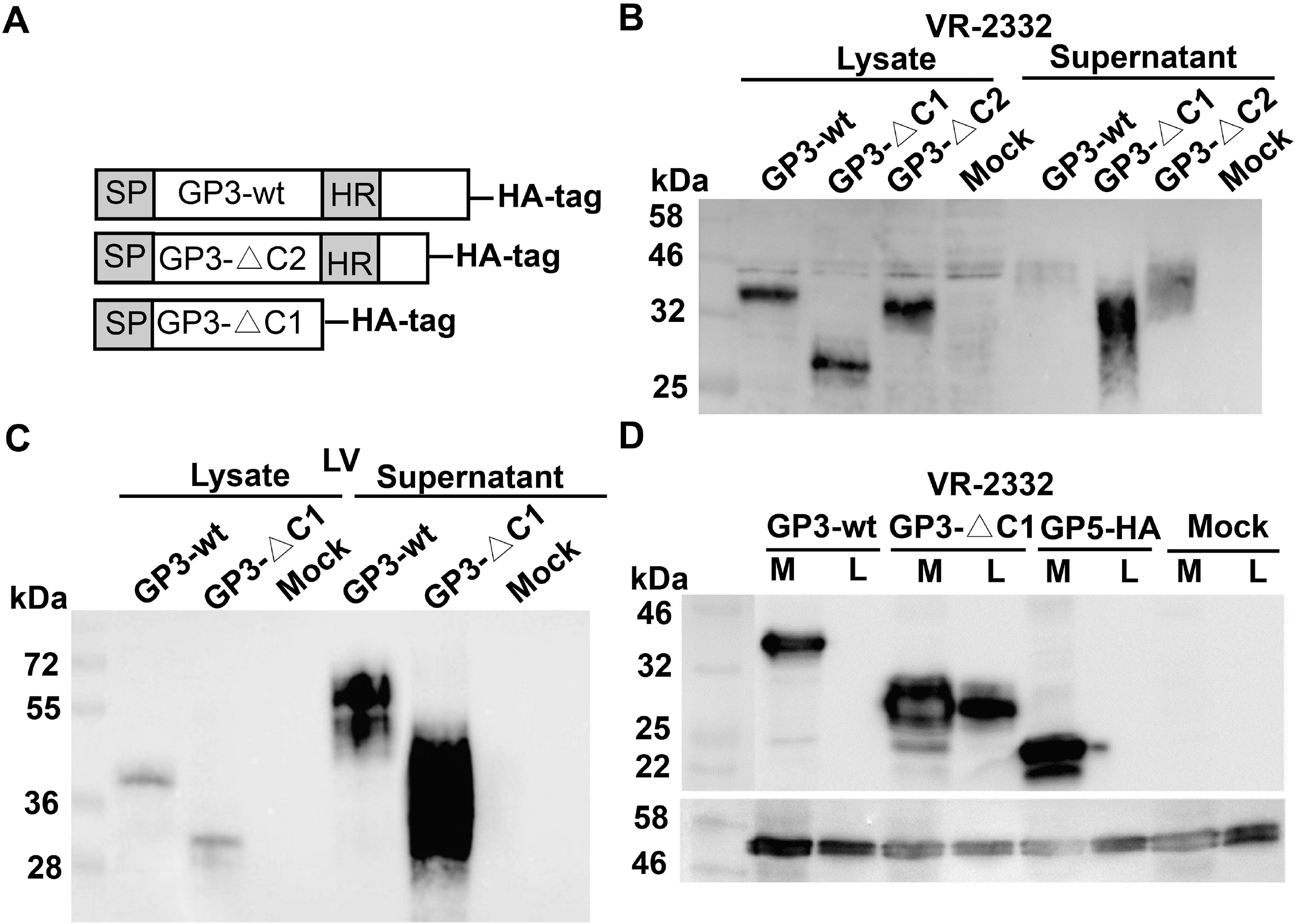
The hydrophobic region at C-terminus of GP3 is a membrane anchor. (A) Scheme of constructs used in these experiments. In GP3-∆C2 the (between PRRSV strains) variable C-terminus (amino acids 230 to 254) was deleted. GP3-∆C1 contains a deletion of The C-terminal part including the hydrophobic region (amino acids 178 to 229 in VR-2332, 172 to 265 in Lelystad virus) SP: signal peptide; HR: hydrophobic region. (B) Release of C-terminal deleted GP3 from VR-2332 into the supernatant of transfected cells [C] Release of C-terminal deleted GP3 from Lelystad virus into the supernatant of transfected cells GP3-HA wild type and mutants GP3-△C1, GP3-△C2 were expressed in BHK-21 cell. 24 hours after transfection the supernatant was removed, cleared and proteins were precipitated with TCA. Cells were washed, trypsinized, pelleted and lysed in denaturing buffer. Samples were subjected to reducing SDS-PAGE and Western blotting with anti-HA antibodies. (D) Release of C-terminal deleted GP3 into the lumen of the ER. 293T cells were transfected with GP3 wild type, mutant GP3-△C1 or (as control) transmembrane protein GP5-HA. 24 hours after transfection cells were gently opened by digitonin, microsomes were prepared by ultracentrifugation, resuspended in buffer and opened by ultrasonication. Membranous (M) and luminal (L) fractions were separated by ultracentrifugation and subjected to Western blotting with antibodies against HA-tag (upper panel) or against calreticulin, a luminal protein of the ER (lower panel).

Western-blotting of cell lysates and supernatants showed that a larger fraction of GP3-∆C1 is now secreted from transfected BHK-cells, whereas the extracellular amount of GP3-∆C2 did only slightly increase relative to GP3-wt (Fig. 4B).Likewise, deletion of the whole C-terminus including the hydrophobic region of GP3 from the PRRSV-1 reference strain Lelystad increased secretion several fold (Fig. 4C).

From this experiment we conclude that deletion of the hydrophobic region enhances secretion of GP3, but it does not prove that GP3 is a membrane protein. One could imagine that the hydrophobic region functions as an ER retention signal and then its removal would allow secretion of a completely soluble protein. We therefore investigated whether intracellular GP3 is membrane-bound and whether the deletion of the hydrophobic region converted intracellular GP3 into a soluble form. Microsomes were prepared from transfected cells, opened by ultrasonication, membranous and luminal fractions were separated by ultracentrifugation and analyzed by Western blotting. The transmembrane protein GP5-HA from PRRSV and calreticulin, a marker for the lumen of the ER, were employed as controls to check the accuracy of the fractionation procedure. GP5-HA, as expected, and also wildtype GP3-HA are only detected in the membrane fraction. GP3-∆C1 is also mainly present in the membrane fraction, but a clear signal was also detected in the luminal fraction. The amount of soluble GP3-∆C1 is probably underestimated, since calreticulin, a soluble protein, also fractionates partly with the membrane, most likely due to the non-quantitative removal of luminal content from the membrane (Fig. 4D). Note also that the soluble fraction of GP3-∆C1 has the same molecular weight as membrane-bound GP3-∆C1. A band with terminal glycosylated carbohydrates that increases the molecular weight is not visible. Thus, once GP3 is released from the membrane it is rapidly secreted from cells as it is more thoroughly been described in the discussion.

Having shown that the hydrophobic region serves as a membrane-anchor of GP3, we asked whether this function can be transferred to another, otherwise soluble protein. We made three constructs in which either the complete C-terminus of GP3, the C-terminus excluding the highly variable region or only the 20 amino acids of the hydrophobic region were fused to the C-terminus of GFP. Fluorescence microcopy of transfected CHO-K1 cells showed that wild-type GFP is present throughout the cell (including the nucleus), whereas all three chimeras localize to an extranuclear, reticular organelle, presumably the ER (Fig. 5A).

**FIG. 5.**
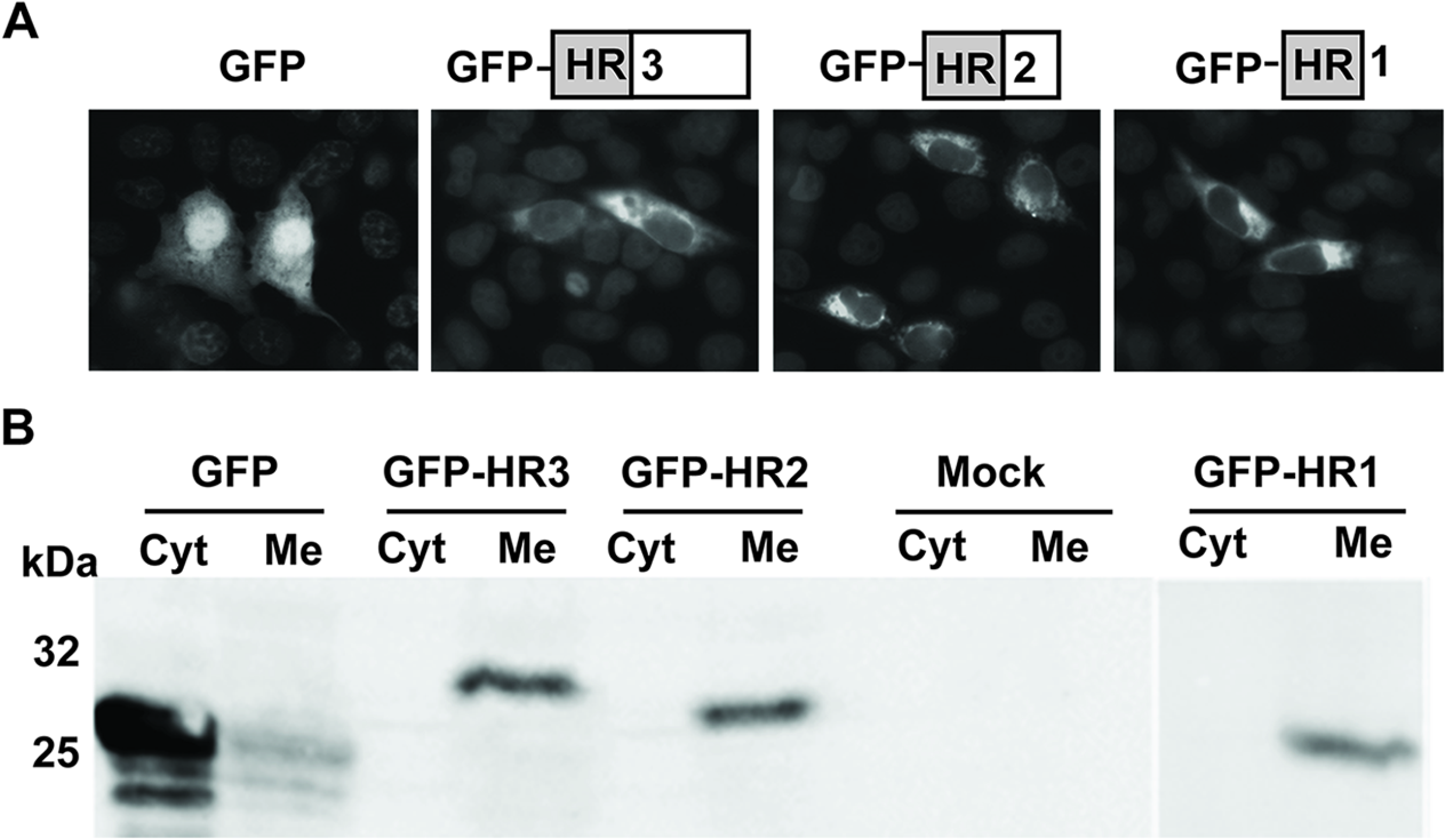
The hydrophobic region of GP3 attaches GFP to membranes. (A) Scheme of GFP-constructs used in these experiments. GFP-HR3 contains the whole C-terminal part of GP3 from VR-2332 (amino acids 178 to 254). GFP-HR2 contains the C-terminal part of GP3 except amino acids 230 to 254, GFP-HR1 contains only the hydrophobic region (amino acids 181 to 200). Localization of GFP and GFP-chimeras in transfected CHO cells visualized with an epifluorescence microscope. (B) CHO cells were transfected with GFP wild type and GFP chimeras, 24 hours after transfection, membrane (Me) and cytosolic (Cyt) fractions were separated by ultracentrifugation and subjected to Western blotting with antibodies against GFP. The SDS-PAGE mobility of molecular weight makers are indicated at the left side of the blot. Mock: untransfected cells.

Fractionation of whole cells into membranes and cytosol (100.000xg pellet and supernatant, respectively) and Western-blotting clearly shows that wild-type GFP is present in the soluble fraction, whereas all three GFP chimeras containing C-terminal GP3 sequences are exclusively membrane-bound (Fig. 5B). Thus, the short hydrophobic region in the C-terminal part of GP3 is sufficient to cause quantitative membrane anchoring of an otherwise soluble protein.

### Exchange of hydrophobic amino acids in the hydrophobic region by alanine increases secretion of GP3

The hydrophobic region is responsible for membrane anchoring of GP3, but the mechanism by which this association is accomplished remains unclear. The bioinformatics program Jpred (http://www.compbio.dundee.ac.uk/jpred/) predicts that the region might form an α-helix (Fig. 6A). Furthermore, this peptide is highly conserved (99 -100%) through GP3 proteins of all PRRSV-1 and PRRSV-2 strains, except at two positions. Position 187 is phenylalanine (98% conservation) in PRRSV-2 strains, but the equivalent position 186 in PRRSV-1 strains (including Lelystad, but not Lena) is leucine (72% conservation). Position 196 is valine in 95% of PRRSV-2 strains, but position 195 is isoleucine in 95% of PRRSV-1 strains (Fig. 6A). The second, slightly variable position is in the middle of an otherwise completely conserved N-glycosylation site NXS. If this site is used the bulky and hydrophilic carbohydrate would certainly not allow binding of GP3 to the ER membrane by hydrophobic forces. Instead, one could imagine that a cellular carbohydrate-binding transmembrane protein, such as Calnexin (45), might anchor GP3. To determine whether the glycosylation site is used, we exchanged the asparagine of NXS to a glutamine generating the mutants N194Q of Lelystad virus and N195Q of VR-2332. Transfection of BHK cells and western blot revealed that wildtype GP3 and the mutants have exactly the same size providing clear evidence that the glycosylation site is not used (Fig. 6B+C), as reported previously for GP3 of the PRRSV-2 FL12 strain (46).

**FIG. 6.**
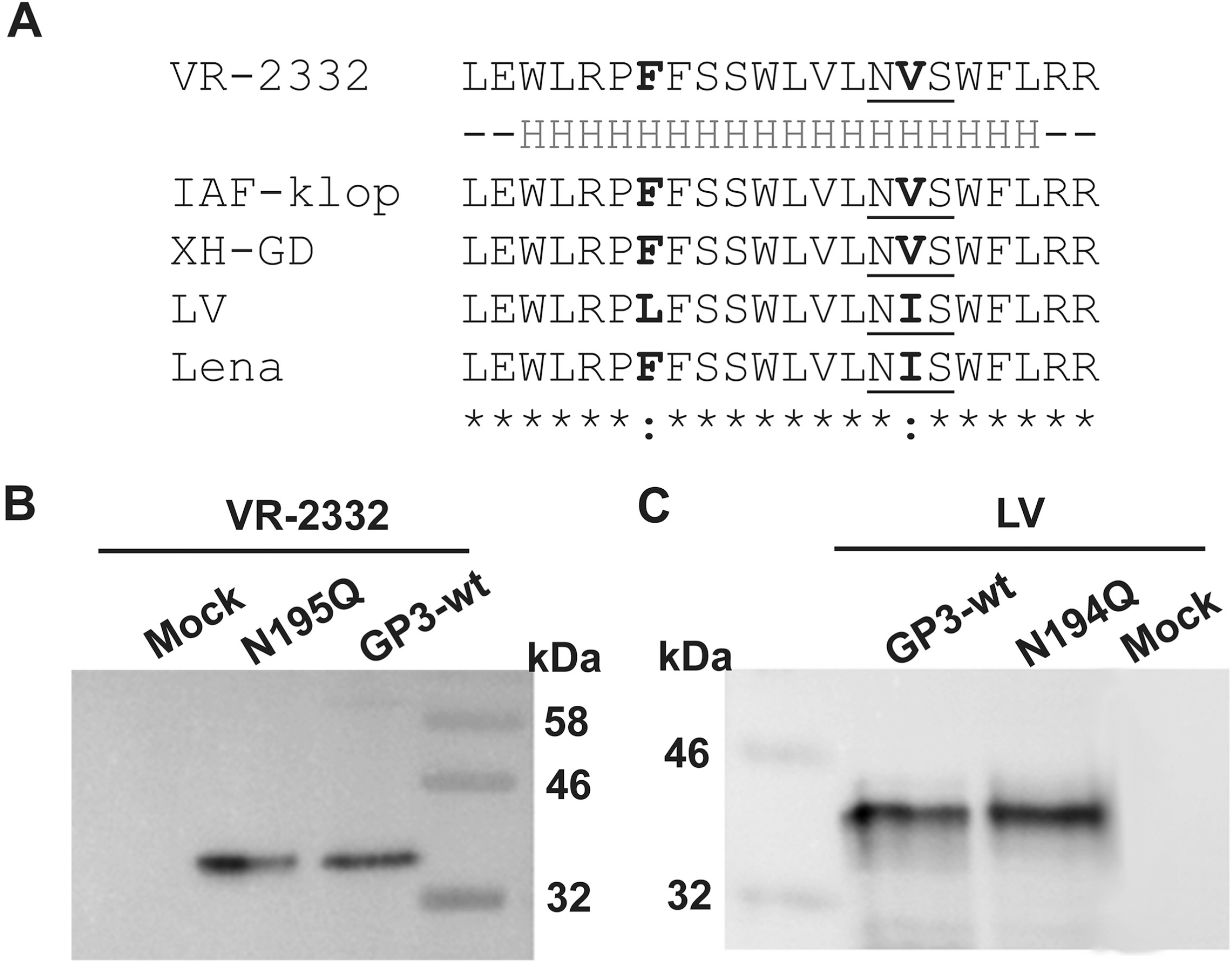
The hydrophobic region is highly conserved and contains an unused glycosylation site. (A) Sequence comparison of the hydrophobic region of GP3. The bioinformatics tool Jpred (http://www.compbio.dundee.ac.uk/jpred/) predicts that amino acids 183-200 of GP3 might from an α-helix (H below the sequence). The region is highly conserved between PRRSV strains. Only two conservative amino acid exchanges (in bold) are present in the used GP3 proteins. The conserved glycosylation site is underlined. (B) and (C) The glycosylation site is not used. Asparagine (N195 in VR-2332 and N194 in Lelystad) of the glycosylation site NXS was exchanged to glutamine (Q) and mutants were expressed in BHK-cells. 24 hours after transfection cells were lysed and samples subjected to SDS-PAGE and Western blotting with anti-HA antibody.

Membrane binding by an amphipathic helix is another mechanism to mediate membrane binding of proteins (47). We therefore employed the bioinformatic tool heliquest (http://heliquest.ipmc.cnrs.fr/) to investigate whether the hydrophobic region might form an aliphatic helix. The distribution of polar versus hydrophobic residues in this putative helix shows that it indeed displays amphipathic characteristics. One surface of the helix contains hydrophobic amino acids, such as leucine, phenylalanine and tryptophan, which favor a lipid environment. The opposite surface also contains hydrophobic amino acids, but they are interrupted by hydrophilic (serine) or even charged ones (arginine), which prefer an aqueous environment (Fig. 7A).

**FIG. 7:**
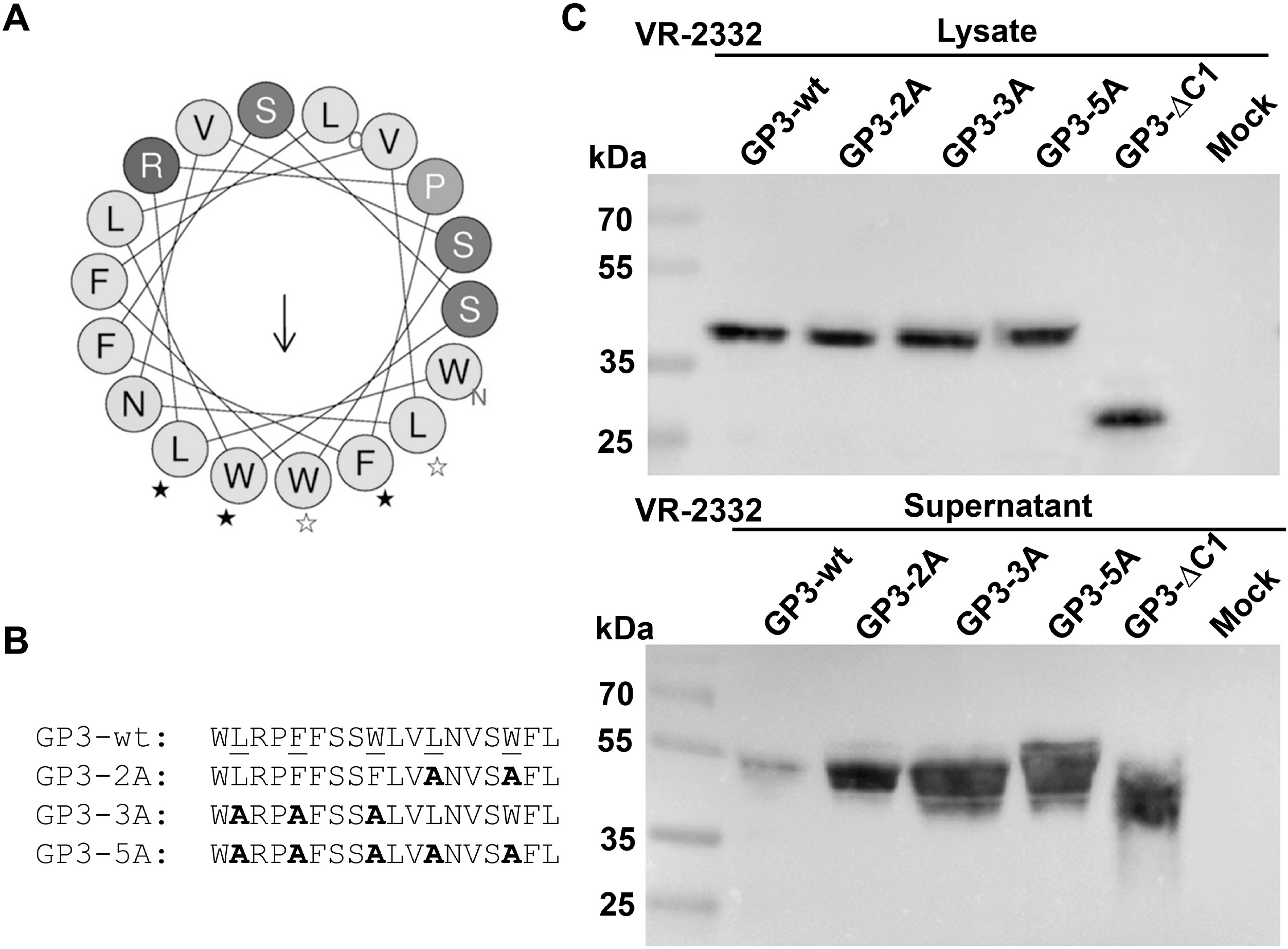
The hydrophobic region might form an amphiphilic helix and exchange of residues in the hydrophobic face enhance secretion of GP3. (A) The bioinformatics tool HeliQuest (http://heliquest.ipmc.cnrs.fr/) predicts that the hydrophobic region of GP3 (amino acids WLRPFFSSWLVLNVSWFL) might form an aliphatic helix with a mean hydrophobicity <H> of 1.131 and a hydrophobic moment <μH> of 0.302. The location of the mutations is indicated by white (mutant 2A) and black asterisks (mutant 3A). (B) Exchange of amino acids in the hydrophobic face by alanines. Underlined amino acids correspond to the amino acids in the lower face of the predicted amphiphilic helix, as indicated by asterisks in (A). (C) BHK-cells cells were transfected with GP3 wild type, and mutant GP3-2A, GP3-3A or GP3-5A. 24 hours after transfection the supernatant was removed, cleared and proteins were precipitated with TCA. Cells were washed, trypsinized, pelleted and lysed in denaturing buffer. Samples were subjected to reducing SDS-PAGE and Western blotting with anti-HA antibodies.

To test the hypothesis that the (presumed) amphipathic helix anchors GP3 to the membrane, we replaced residues in its hydrophobic face by alanine. In GP3-2A leucine 194 and tryptophan 198 were replaced, the mutant GP3-3A contains exchanges of leucine 184, phenylalanine 187 and tryptophan 189. GP3-5A combines all five exchanges such that the predicted membrane insertion face of the helix is completely replaced by alanine (Fig. 7B). GP3-HA wildtype and the mutants were expressed in BHK-21 cells, and the distribution of GP3 protein in cells and supernatants was analyzed. The Western-blot clearly shows that secretion of all mutants is greatly enhanced to a similar extent as removal of the complete C-terminal part of GP3 in the mutant GP3-∆C1 (Fig. 7C).

### Substitution of hydrophilic by hydrophobic amino acids in the amphiphilic region prevents secretion of GP3

To further substantiate that the hydrophobic region determines the strength of membrane anchoring of GP3 we replaced hydrophilic with more hydrophobic residues. The mutant GP3-4H contains an exchange of arginine 185 and proline 186 by leucine plus a replacement of serine 189 and 190 by phenylalanine (black asterisks in Fig, 8A). The exchanged amino acids are solely located in the hydrophilic face of the predicted amphiphilic helix. In the mutant GP3-3H also amino acids in the hydrophobic face were exchanged; asparagine 195 was replaced by a serine and serine 197 plus tryptophan 198 were replaced by isoleucines (white asterisks in Fig. 8A). GP3-7H combines all seven exchanges (Fig. 8B). When these three mutations were created in GP3 of the strain XH-GD secretion of the protein into the cellular supernatant was completely prevented indicating that the protein is now firmly membrane-anchored (Fig. 8C).

**FIG. 8.**
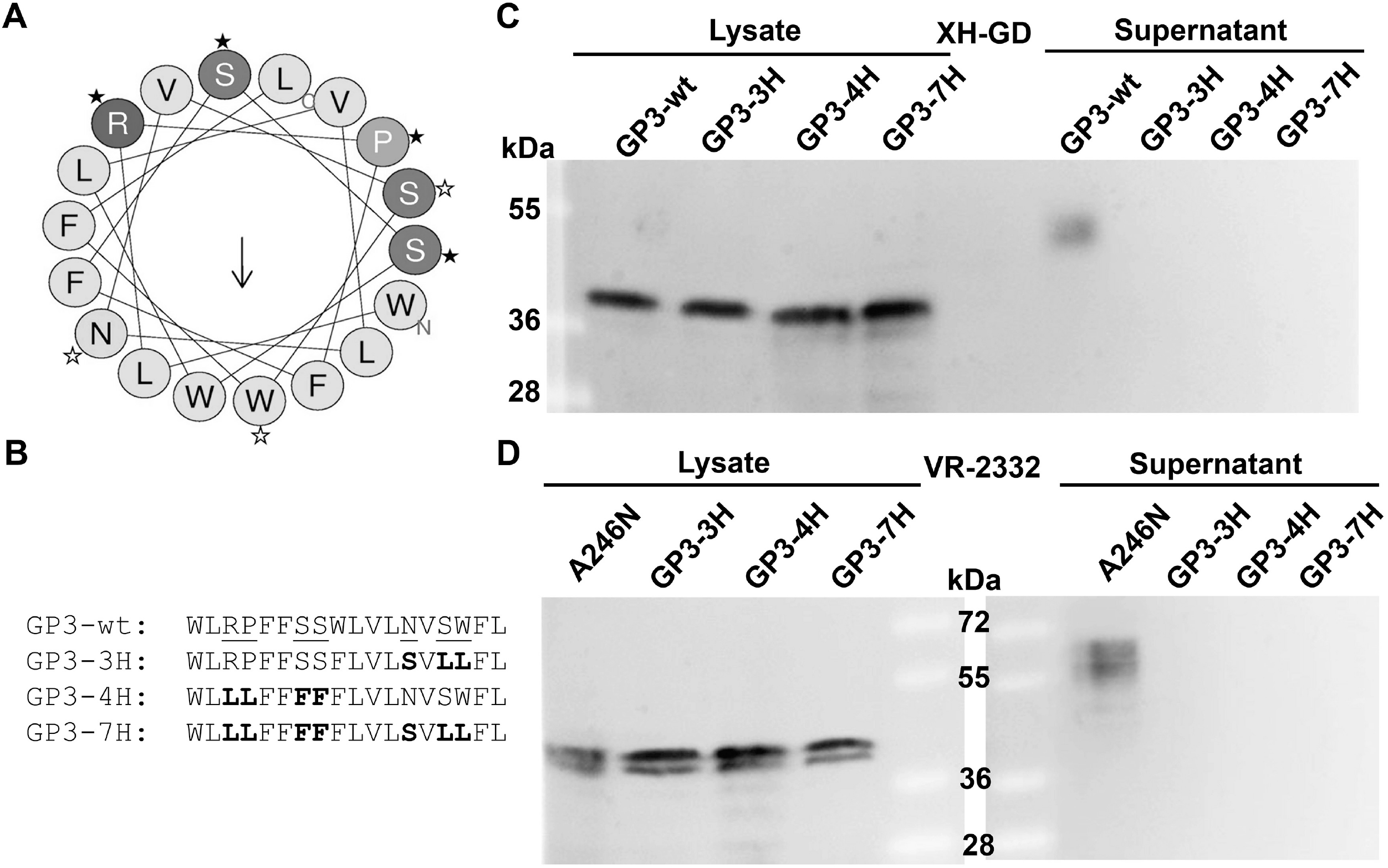
Exchange of residues in the hydrophilic face by hydrophobic amino acids prevents secretion of GP3. (A) Predicted aliphatic helix of GP3 with location of mutations indicated by white (mutant 3H) and black asterisks (mutant 4H). (B) Exchange of hydrophilic amino acids by the more hydrophobic amino acids (leucine (L), phenylalanine (F) and serine (S). (C) BHK-cells cells were transfected with mutant GP3-wt and GP3-3H, GP3-4H or GP3-7H of XH-GD. (D) BHK-cells cells were transfected with mutant GP3-A246N (which contains an additional glycosylation site the C-terminal region) and mutant GP3-3H, GP3-4H or GP3-7H (D), which also contain this additional glycosylation site. 24 hours after transfection the supernatant was removed, cleared and proteins were precipitated with TCA. Cells were washed, trypsinized, pelleted and lysed in denaturing buffer. Samples were subjected to reducing SDS-PAGE and Western blotting with anti-HA antibodies.

In principle, this could be achieved by two mechanisms. Replacing hydrophilic by hydrophobic amino acids creates an uninterrupted stretch of 18 hydrophobic residues, which is long enough to span the ER membrane (48). This would create a transmembrane protein with the C-terminal region exposed to the cytosol. Alternatively, despite the mutations, the C-terminus (including the hydrophobic region) is completely translocated into the lumen of the ER. To distinguish between both possibilities we also tested the three mutants in the context of the GP3 mutant A264N of VR-2332, which contains the additional (and used) glycosylation site in the C-terminal part (Fig. 3C). Similar to GP3 from XH-GD the mutations prevent secretion of GP3 into the cellular supernatant. Comparing the intracellular band pattern revealed that in all three mutants the additional glycosylation site is used to the same extent as in GP3 without mutations in its hydrophobic region (Fig. 8D). Thus, the C-terminal part is translocated into the ER lumen; the protein does not adopt a transmembrane topology. In sum, we obtained clear evidence that the hydrophobic region determines membrane anchoring of GP3 probably by forming an amphiphilic helix, but a direct structural analysis is required to prove the later assumption.

### The between PRRSV-1 and PRSV-2 genotypes variable C-terminus determines the amount of secreted GP3

Having established that the highly conserved hydrophobic region is the main membrane anchor of GP3 we asked whether other between PRRSV-1 and PRRSV-2 strains variable features modulate how much of the wild type protein is secreted. A Kyte–Doolittle hydropathy plot revealed a remarkable difference in the biophysical properties of the C-terminal part of GP3; it is rather hydrophobic in GP3 from the PRRSV-2 strain XH-GD, but strongly hydrophilic in the PRRSV-1 strain Leystad (Fig. 9A). We hypothesized that the different hydrophobicity in the C-terminus might affect the strength of membrane attachment of GP3s. We therefore deleted the C-terminus from GP3 from both PRRSV strains. This deletion considerably enhanced secretion of GP3 from XH-GD (Fig. 9C), but had no obvious effect on secretion of GP3 from Lelystad (Fig. 9B) which is already secreted in higher amounts compared to GP3 from XH-GD (Fig. 1A). Exchanging the C-terminus between both GP3 proteins completely reversed this secretion behavior. GP3 from Lelystad with the C-terminal domain of XH-GD is secreted in lower amounts than the corresponding wild-type protein (Fig. 9D), whereas GP3 from XH-GD with the C-terminal domain of Lelystad is secreted in higher amounts (Fig. 9E). In sum, the between PRRSV-1 and PRRSV-2 strains variable C-terminal domain modulates how much of GP3 is secreted.

**FIG.9. The.**
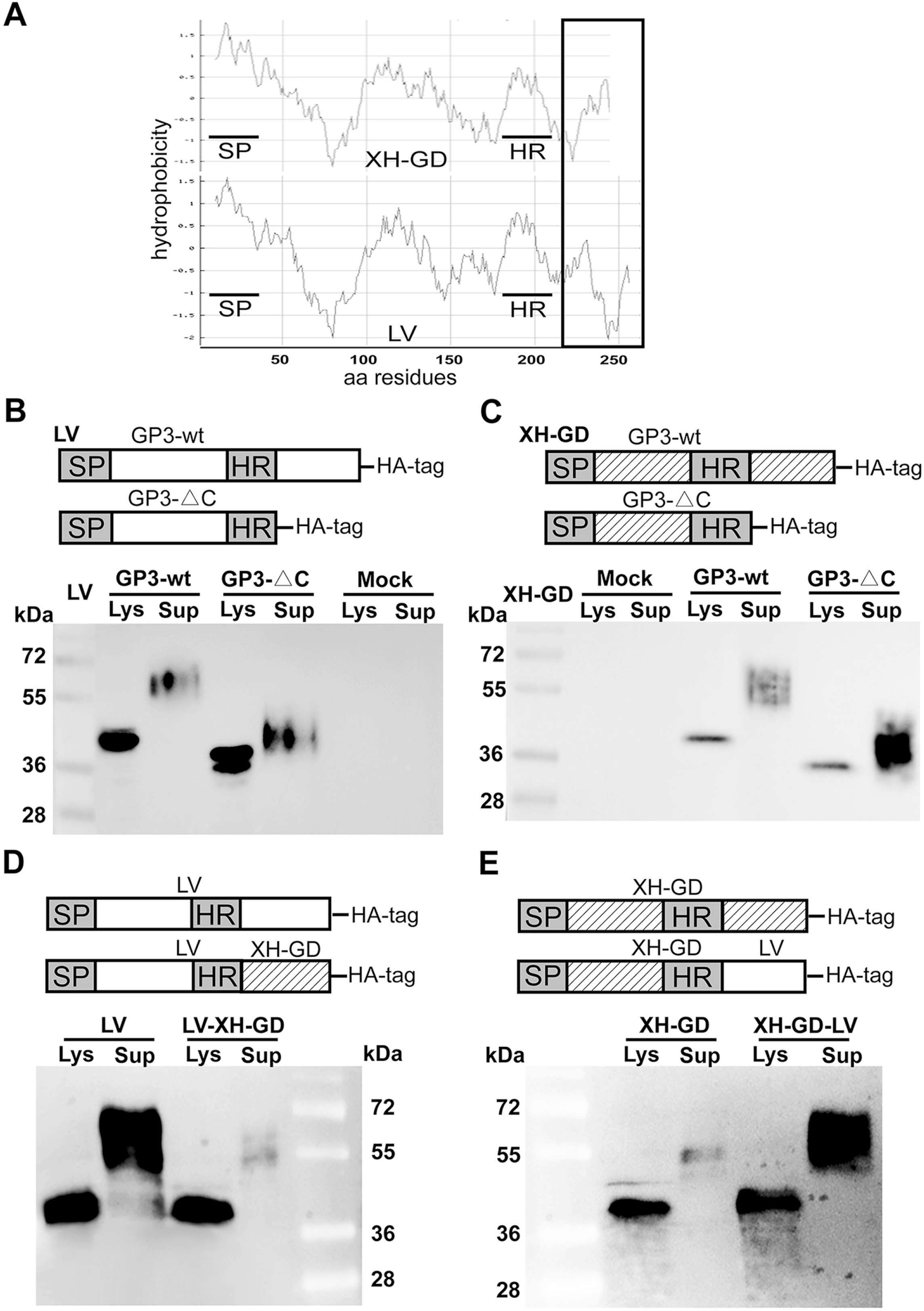
variable C-terminus modulates secretion of GP3. (A) Kyte-Doolittle hydropathy plot of GP3 from Lelystad (LV) and XH-GD PRRSV strains. y-axis: hydropathy score, x-axis: amino acid position. SP: signal peptide, HR: hydrophobic region. The black rectangle highlights the variable C-terminal region which is hydrophilic in GP3 from Lelystad, but hydrophobic in Gp3 from XH-GD strain. (B) and (C) Deletion of the between strains variable C-terminal domain from GP3. In GP3-ΔC from Lelystad (LV) amino acids 204-265 and in GP3-ΔC from XH-GD amino acids 205-254 were deleted. Constructs of Lelystad (LV) (B) and XH-GD (C) were expressed together with the corresponding wild type proteins in BHK-21 cell. (D) and (E) Exchange of the variable C-terminal domain of GP3 between PRRSV-1 and PRRSV-2 strains. (D) Creating the chimera LV-XH-GD: amino acids 1-203 of GP3 from Lelystad were fused to amino acids 205-254 of GP3 from XH-GD. (E) Creating the chimera XH-GD-LV: amino acids 1-204 of GP3 from XH-GD were fused to amino acids 204-265 of GP3 from LV. The chimeras contain an HA-tag fused to their C-termini. Constructs of Lelystad (LV) (D) and XH-GD (E) were expressed together with the corresponding wild type proteins in BHK-21 cell. 24 hours after transfection the supernatant was removed, cleared and proteins were precipitated with TCA. Cells were washed, trypsinized, pelleted and lysed in denaturing buffer. Samples were subjected to reducing SDS-PAGE and Western blotting with anti-HA antibodies.

### Mutation of the hydrophobic region inhibits replication of PRRSV

Finally, we analyze whether mutations in the hydrophobic region of GP3 affect virus replication. We used a reverse genetics system based on a full-length cDNA clone of the Chinese PRRSV-2 strain XH-GD cloned into a plasmid downstream of a CMV promotor. We created the same three mutants that caused enhanced secretion of GP3-HA from VR-2332, but this was not possible without altering also amino acids at the N-terminus of the overlapping GP4 gene. However, the changes are located entirely in the (variable) signal peptide of GP4 and are predicted by SignalP (http://www.cbs.dtu.dk/services/SignalP/) not to change its cleavability and hence the functionality of the mature GP4 protein (see “materials and methods” for details).

Full-length cDNA was transfected into CHO-K1 cells (which lack the receptors for cell entry of PRRSV) and 48 hours later the supernatant was used to infect MARC-145 cells, which support virus replication. Immunofluorescence using antibodies against GP5 revealed that a similar percentage of cells synthesize the viral glycoprotein, i. e. the transfection efficiency was comparable with wildtype or mutant cDNA (Fig. 10A). However, infected cells show a fluorescence signal only for wildtype virus, not for any of the three mutants. Not any immunofluorescence signal was detected if the passage one supernatant was subjected to two further passages in MARC-145 cells (not shown). Likewise, two further attempts to rescue infectious mutant virus (including transfection of BHK-21cells) also failed (not shown). We conclude that PRRSV mutants are able to synthesize viral proteins in transfected cells, but cells apparently do not release infectious virus particles.

**FIG. 10.**
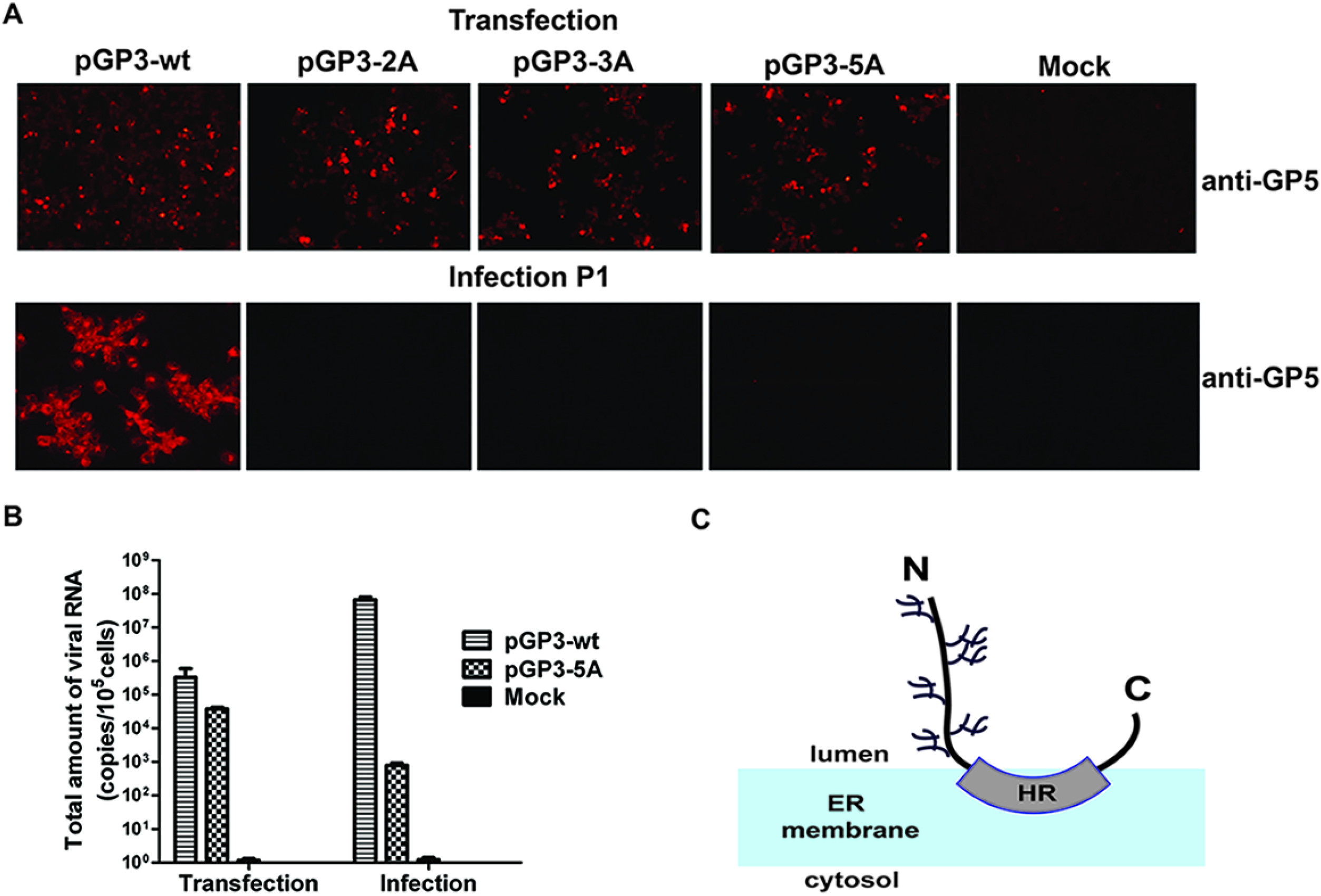
Mutations in the predicted amphipathic helix prevent virus replication. (A) Immunofluorescence of cells transfected (upper panel) or infected (lower panel) with full-length clone of the PRRSV strain XH-GD, either wildtype (pGP3-wt) or the three alanine mutants pGP3-2A, pGP3-3A or pGP3-5A having the same mutations as shown in Fig. 7A and 7B. Upper panel: CHO-K1 cells were transfected, ~48 h later the supernatant was removed and permeabilized cells were stained with anti-GP5 monoclonal antibody and Alex-568 anti-mouse secondary antibody. Lower panel: MARC-145 cells were infected with the supernatant and 24 h post infection stained with anti-GP5 monoclonal antibody and Alex-568 anti-mouse secondary antibody. (B) Quantification of viral genomes (pGP3-wt and pGP3-5A) released into the supernatant of transfected or infected cells. 24 hours post-transfection or postinfection, viral RNA was isolated from cleared culture supernatants and reverse transcribed into cDNA. Full-length plasmid of infectious clone pGP3-wt was as used as a standard to estimate the number of gene copies. Results from two independent transfections or infections are shown as number of viral RNA (mean including standard deviation) released into the supernatant of 1×10^5^ transfected or infected cells. (C) Model of GP3. HR: hydrophobic region which attaches GP3 to the membrane. N and C indicate the N – and C-terminus of the protein. The branches are the carbohydrates attached to GP3 of VR-2332. See Figs. 1B-E and 2A for number of carbohydrates attached to GP3 proteins of other PRRSV strains.

For both PRRSV and EAV it was reported that blockade of GP3 expression in an infectious clone prevents rescue of infectious virus, but virus-like particles (lacking GP2, GP3 and GP4) were released into the supernatant of transfected cells (11, 12). To analyze whether cells transfected with pGP3-5A cDNA having five amino acid substitutions in the helix exhibit the same phenotype we used qRT-PCR to estimate the number of viral genomes released into the supernatant of transfected CHO-K1 and infected MARC-145 cells. Comparable gene copy numbers were detected in the supernatant of cells transfected with wt cDNA (5×10^5^) and pGP3-5A cDNA (8×10^4^) indicating that the mutation does not prevent release of particles. When this supernatant was used to infect MARC-145 cells, 5×10^8^ RNA gene copies of wild type virus were detected 48 hours later (indicating that the input virus multiplied ~1000fold), but very little (10^3^) for the pGP3-5A mutant. Since the number of mutant genomes was greatly reduced (8×10^4^ →10^3^), the viral genomes still present in the supernatant of “infected” cells probably originate from the input virus. In sum, we conclude that the apolar face of the presumed amphipathic helix of GP3 is not essential for budding, but for virus entry.

## DISCUSSION

In this study we investigated secretion, membrane-anchoring and the membrane topology of GP3 from PRRSV-1 and PRRSV-2 strains. We propose that GP3 exhibits an unusual hairpin-like membrane topology (Fig. 10C). We show that the C-terminal part of GP3 is exposed to the lumen of the ER since it is protected against proteolytic digestion in perforated cells (Fig. 3B) and since inserted N-glycosylation sites are efficiently used (Fig. 3C and 3D). This excludes that GP3 is a classical type I membrane protein containing a transmembrane region and a cytosolic tail. Nevertheless, the hydrophobic region serves as a membrane anchor. Its deletion enhances secretion of GP3 from transfected cells by several orders of magnitude (Fig. 4B+C). The bioinformatics tool heliquest predicts that the short hydrophobic region forms an amphipathic helix (Fig. 7A). Amphipathic helices can adopt an orientation parallel to the membrane plane; hydrophobic residues with long side chains insert between fatty acyl-chains of the lipids, whereas polar residues face their polar heads (47). Exchange of only two or three hydrophobic amino acids in the hydrophobic face of the presumed helix by alanine (which are also hydrophobic, but have only a short methylene group as side chain) increases secretion of GP3 to the same extent as removal of the complete C-terminal part of GP3 (Fig. 7C). Likewise, replacement of three or four hydrophilic by hydrophobic residues prevents secretion of GP3 (Fig. 8) indicating that the hydrophobic properties of amino acids in this region determine the strength of membrane binding.

Finally, we could not generate infectious PRSSV virus having the same exchanges in the hydrophobic face of the helix of GP3 suggesting that the domain is essential for a complete replication cycle. However, viral genomes of the mutants (and thus probably virus particles) were released into the supernatant of transfected cells (Fig. 10). Thus, these mutants show an identical phenotype as mutants of the equine arteritis virus where expression of the GP3 gene was blocked (11, 12), reinforcing the assumption that a functional GP3 is not essential for budding, but for virus entry into target cells.

A multitude of amphipathic helices that interact with membranes have recently been characterized in cellular proteins. Their interactions are mostly reversible and restricted to certain cellular membranes in order to allow the respective protein to fulfil a specific cellular function. The binding specificity is often determined by interactions with certain lipids, for example basic amino acids in the helix interact with negatively charged lipids, such as phosphatidylserine or phosphatidylinositols (47).

Amphipathic helices are characterized by two physico-chemical parameters, the hydrophobic moment (<μH>) and the average hydrophobicity (<H>), which are calculated by heliquest. The hydrophobic moment tells us if a sequence, when considered as helical, exhibits one hydrophobic face and one polar face whereas the hydrophobicity describes the avidity of the helix for lipids (49). Calculating these parameters for the presumed helix of GP3 reveals that its hydrophobic moment is compatible with being amphipathic (0.302) and the average hydrophobicity is high (GP3: 1.131) suggesting a more stable membrane binding than transiently membrane-bound helices. Indeed, the hydrophobic region of GP3 confers complete membrane binding when fused to GFP, an otherwise soluble protein (Fig. 5). Nevertheless, its membrane anchoring ability is most likely lower than that of a transmembrane region. Once inserted into the membrane, transmembrane regions are permanently anchored; release (without damaging the bilayer) is not possible. Although shedding of viral type 1 membrane glycoproteins have been described, but they are often generated by proteolytic removal of the transmembrane region (42). However, this is not the case for GP3 since deglycosylated GP3 present inside cells and in the supernatant have identical SDS-PAGE mobility (Fig. 1).

Our results are best interpreted if we assume that membrane-bound and soluble GP3 exist in equilibrium. Wildtype GP3 is mostly membrane-bound in the ER, but some GP3 molecules might detach from the membrane and escape from this organelle by inclusion into the lumen of COPII vesicles, the carriers for transport from the ER to the Golgi. In the Golgi, all carbohydrates are processed to an Endo-H resistant, terminally glycosylated form that is secreted from the cells (Fig. 1F). Identical results have been reported previously for GP3 of the IAF-Klop strain (30).

Note, however, that GP3 with Endo-H resistant carbohydrates and a higher molecular weight was not observed in cell lysates (Fig. 1+4) suggesting that transport of GP3 through the Golgi is very fast. This is in line with the general concept that the rate-limiting step for protein transport along the exocytic pathway is exit from the ER; especially small proteins revealed very short half times for Golgi-transit of only 10 minutes (50). Thus, under steady state conditions (transfected cells) most GP3 molecules are either in the ER or already secreted; only a very small fraction of Endo-H resistant GP3 is present in the Golgi and this is therefore not detectable by Western blotting.

This very fast processing also suggests that soluble GP3 once transported to the Golgi does not rebind to its membranes; i. e. the equilibrium is now shifted to the soluble form. The reason might be that the physicochemical nature of the lipid bilayer changes along the exocytic pathway. The cholesterol content increases from ~10% to 45%, which causes tighter packaging of lipids and this might disfavor insertion of hydrophobic amino acid side chains. In addition, the asymmetric lipid distribution characteristic of the plasma membrane begins to build up in the Golgi, e. g. negatively charged lipids are translocated from the luminal to the cytoplasmic side of the membrane (51). These lipids might enhance binding of GP3´s helix to the ER membrane (a positively charged arginine is present in the helix), but their number might be reduced in the luminal part of Golgi membranes. However, detailed experiments with purified GP3 and artificial membranes are required to elucidate the lipid specificity of the helix and the amino acids essential for binding.

Exchange of hydrophobic residues in the apolar face of the helix by alanine shifts the membrane binding equilibrium of GP3 to the soluble version. Heliquest predicts that the average hydrophobicity of the helix is only moderately reduced in the mutants (wt: 1.131; mutant 2A: 0.946; 3A: 0.863; 5A: 0.678), but the hydrophobic moment is severely diminished in each mutant (wt: 0.302; 2A: 0.155, 3A: 0.068, 5A: 0.107) suggesting that this part of the molecule is no longer able to form an amphipathic helix. As a result mutant GP3s bind to membranes only poorly and a much higher fraction is secreted from cells (Fig. 7).

In contrast, exchange of hydrophilic by hydrophobic residues in the polar face of the helix shifts the membrane binding equilibrium of GP3 to the membrane anchored form (Fig. 8). Accordingly, the mutations successively increase the hydrophobicity of the hydrophobic region (wt: 1.131, mutant 3H: 1.228; 4H: 1.539; 7H: 1.636) and also decreased the hydrophobic moment (wt: 0.302; 3H: 0.271, 4H: 0.076, 7H: 0.098).

Interestingly, the hydrophobic region also contains a completely conserved glycosylation site (N195 in VR-2332), which is not used (Fig. 6), presumably because membrane binding prevents access of the oligosaccharyltransferase to this site. The glycosylation site might become accessible for glycosylation in mutants with diminished membrane binding affinity since both intracellular and secreted forms of GP3-5A exhibit a higher molecular weight compared to wildtype GP3 (Fig. 7C). Other Western-blots having better separation revealed an additional band with a slightly higher molecular weight for each mutant consistent with the conclusion that a fraction of ~10% of GP3-2A and GP3-3A and ~50% of GP3-5A are glycosylated at N 195 (not shown).

Exchange of the hydrophobic residues by alanine in the presumed amphipathic helix of GP3 prevented release of infectious virus particles from transfected cells (Fig. 10). We assume that a (mostly) soluble version of GP3 does not form a functional GP2/3/4 complex, which is a prerequisite for virus entry (11, 12). We could not test whether GP3 (or GP2 or GP4) are incorporated into virus particles since antibodies against the minor glycoproteins of the XH-GD strain are not available and insertion of an HA-tag into a full-length clone of PRRSV is not possible since the C-terminus of GP3 overlaps with the N-terminus of GP4.

The essential nature of the hydrophobic region is also corroborated by the notion that it is highly conserved between GP3 proteins from all PRRSV-1 and -2 strains (Fig. 3A, Fig. 6A). A predicted amphipathic helix with a very similar sequence is also present in GP3 of LDV (**<u>YI</u>**RPLFSSWLVLNVSY**<u>F</u>**L, three conservative exchanges to PRRSV VR-2332 are underlined, <μH>: 0.215; <H>: 0.988) and it was reported that a fraction of the protein is secreted from cells and that it can be extracted from membranes without detergent suggesting that it is a peripheral membrane protein (29). GP3 from EAV also adopts a hairpin-like topology and is membrane-anchored by a C-terminal hydrophobic domain (33). This region (RPTLICWFALLLVHFLPMPRCRGS) exhibits no sequence homology to the presumed helices of PRRSV and LDV, but heliquest predicts that it forms an amphipathic helix with similar biophysical properties (<μH>: 0.258; <H>: 1.181 for underlined amino acids) than the helices of PRRSV and LDV. Similar to PRRSV,membrane-anchoring of GP3 of EAV is also essential for virus replication since infectious mutants with a stop codon inserted into this region could not be generated (34). In sum, although experimentally not analyzed for each protein, GP3 proteins from PRRSV, LDV and EAV might have the same hairpin-like membrane topology and the same mechanism of membrane-anchoring, but the presumed amphipathic helix is formed by different amino acids in the genera Rodartevirus (LDV and PRRSV) and Equartevirus (EAV).

Although they contain an (almost) identical hydrophobic region, a larger fraction of GP3s from the PRRSV-1 strains is secreted from transfected cells compared to GP3s from the PRRSV-2 strains (Fig. 1). The one (Lena) or two (Lelystad) conservative exchanges in the helix reduced its physico-chemical parameters only slightly (Lena: H: 1.163; μH: 0.273, Lelystad: H: 1.158; μH: 0.269). Instead we have shown that the variable C-terminus modulates how much of GP3 is secreted. The C-terminal part is strongly hydrophilic in the PRRSV-1 strain Lelystad, but rather hydrophobic in the PRRSV-2 strain XH-GD (Fig. 9A). Exchanging the C-terminus between both GP3 proteins completely reversed this secretion behavior. GP3 from Lelystad with the C-terminal domain of XH-GD is secreted in lower amounts than the corresponding wild-type protein (Fig. 9D), whereas GP3 from XH-GD with the C-terminal domain of Lelystad is secreted in higher amounts (Fig. 9E).

Note, however, that the C-terminus is the most variable part of GP3 (Fig. 3A, the sequence also reveals insertions or deletions) and is thus remains to be shown whether increased GP3 secretion is a general feature of all PRRSV-1 strains. However, the secreted form of GP3 is likely to be a folded (and hence functional) protein, since it passed the quality control system of the ER (45). Furthermore, since a large fraction of secreted (but not intracellular) GP3 is a disulfide-linked dimer, at least one intermolecular disulfide bond is formed which would be unlikely to happen in a misfolded protein (Fig. 1G).

The only other viral structural glycoprotein having a similar membrane anchor is the E^rns^ glycoprotein of Pestiviruses, an unspecific RNAse that suppresses the cellular innate immune response and is involved in the establishment of persistent infection. However, the amphiphilic helix of E^rns^ is longer (60 amino acids) and located at the extreme C-terminus of the protein. Another similarity to GP3 is the dual nature of E^rns^. E^rns^ is an essential part of the viral envelope, but is also secreted into the extracellular medium and found in the serum of infected animals. This secreted form is hypothesized to relate to its role as a virulence factor (52–54).

We speculate that secreted GP3 might also play a role during PRRSV infection of pigs, for example as a decoy that serves to distract antibodies away from virus particles. Indeed, antibodies against various regions of the GP3 protein were elicited in the majority of experimentally infected piglets, but most of them had no or only little neutralizing activity (19–25).

## ACKNOWLEDGEMENTS

This work was supported by the German Research Foundation (DFG, grant number VE 141/13-1); Minze Zhang is recipient of a PhD fellowship from the China Scholarship Council (CSC). Fangkun Wang´s sabbatical at the Free University Berlin was supported by the “growth of young teachers program” of the Shandong province. The funders had no role in study design, data collection and interpretation, or the decision to submit the work for publication.

